# BAM15 treats mouse sepsis and sepsis-AKI, linking circulating mitochondrial DNA and tubule reactive oxygen species

**DOI:** 10.1101/2021.07.06.451287

**Authors:** Naoko Tsuji, Takayuki Tsuji, Tetsushi Yamashita, Xuzhen Hu, Peter S.T. Yuen, Robert A. Star

## Abstract

The pathogenesis of sepsis is complex and heterogeneous; hence, a precision medicine strategy may be required. Acute kidney injury (AKI) following sepsis portends higher mortality. Overproduction of mitochondrial reactive oxygen species (mtROS) is a potential mediator of sepsis and sepsis-induced AKI. BAM15 is a chemical uncoupler that dissipates the mitochondrial proton gradient without generating mtROS, and improves experimental renal ischemic injury. We injected BAM15 into mice at 0 or 6 hours after cecal ligation and puncture (CLP) treated with fluids and antibiotics. BAM15 reduced mortality, even when started at 6 hours, when mice were ill, and reduced kidney damage but did not affect other organs. Serial plasma and urinary levels of mitochondrial DNA (mtDNA) were increased following CLP, and decreased after BAM15 (at 0 and at 6 hours). *In vitro* BAM15 prevented mtROS overproduction and mtDNA release from septic kidney tubule cells; mtROS generation correlated with mtDNA release. BAM15 also promotes mitochondrial biogenesis signaling. We conclude that BAM15 is an effective preventive and therapeutic candidate in experimental sepsis, and that BAM15 and mtDNA are mechanistically linked via mtROS, which may form a drug-companion diagnostic pair to improve precision medicine approaches to diagnosing and treating clinical sepsis.

## Introduction

Sepsis is caused by severe infection and progresses to multiple organ failure including acute kidney injury (AKI). Globally, 48.9 million cases of sepsis are reported world-wide and 11 million sepsis-related deaths were reported, representing 20% of all global deaths (1). One third of patients with sepsis will develop AKI (sepsis-AKI), with a considerable increase in mortality (2). Sepsis accounts for 40% of AKI among critically ill patients (3) and sepsis-AKI has higher mortality than other forms of AKI (1). Even with infectious source control, antibiotics, and renal replacement therapy (RRT), sepsis-AKI portends a very high mortality. There is no effective drug to prevent or treat human sepsis or sepsis-AKI, in part because of multifactorial pathophysiologies that differ amongst patients with clinical sepsis. Clinical trials are difficult to perform because it is unclear who to treat, with what drug, and at what dose. A targeted precision medicine approach has been suggested (4, 5), which would be enhanced by co-development of drugs and co-developed biomarkers, as in cancer (6).

Mitochondrial DNA, recognized recently as a damage-associated molecular pattern (DAMP), is released from injured tissues and circulating, activated immune cells (7). We and others have shown that cell-free (cf) mtDNA is released into the circulation during the early phase of sepsis (8, 9) and also in patients with severe SARS-COV2 infection (10–12). In sepsis, mtDNA contributed to AKI with overproduction of renal mitochondrial superoxide via Toll-like receptor 9 (TLR9) (8). Thus, cfmtDNA might be both a biomarker and a therapeutic target of disease severity in sepsis-AKI. However, cfmtDNA has not been evaluated as a therapeutic efficacy biomarker in sepsis or sepsis-AKI.

BAM15 [(2-fluorophenyl)-(13-e]pyrazin-5-yl))amine] is a mitochondrial uncoupler that can protect the kidney from cold storage-induced damage (14) and ischemic injury (13). Mitochondrial uncouplers are small chemical compounds that transport protons from the mitochondrial inner membrane space into the mitochondrial matrix independent of ATP synthase, and thus uncouple nutrient metabolism from ATP generation. Mitochondrial uncouplers paradoxically decrease superoxide production by increasing the speed of electron transfer through the electron transport chain, thus decreasing electron dwell time in the electron transport chain (15–17). BAM15 has less cytotoxicity and fewer off-target effects on plasma membrane depolarization compared to other uncouplers (13). The therapeutic potential of mitochondrial uncouplers has been investigated in the treatment of obesity (18), type 2 diabetes (19), and ischemic reperfusion injury-induced AKI (13), but not in sepsis or sepsis-AKI. In endotoxin- or cisplatin-induced AKI, mitochondrial and renal function can be restored if mitochondrial damage is prevented by decreasing excess mtROS and increasing mitochondrial biogenesis signaling, which is inhibited during AKI (20, 21). Therefore, we investigated the effect of BAM15 as a mitochondria-targeted drug for sepsis-AKI, and evaluated mtDNA as an efficacy biomarker for BAM15 treatment.

## Results

### BAM15 increased sepsis survival and reduced sepsis-AKI, even with delayed treatment

We first evaluated the impact of BAM15 on clinically relevant outcomes of sepsis and sepsis-AKI. We performed a survival study using a fluid- and antibiotic-treated cecal ligation and puncture (CLP) model of mouse sepsis (22) (23). Early treatment with BAM15 (5 mg/kg i.p.) at the time of CLP surgery substantially increased survival (survival at 7 days, CLP+Vehicle: 25% vs CLP+BAM: 75%, p<0.05) (Figure 1A). Early treatment with BAM15 also improved kidney dysfunction at 18 hours after CLP (serum creatinine: CLP+vehicle vs CLP+BAM15: 0.43± 0.05 vs 0.12±0.02 mg/dl, p<0.05; BUN, CLP+vehicle: 103.8 ± 16.2 mg/dl vs CLP+BAM15: 53.4 ± 11.5 mg/dl, p<0.05), without a significant effect on other organ damage indices (Figure 1B-C). The benefit of BAM15 pre-treatment was also observed in female mice (Supplemental Figure 1A-C). Delayed treatment with BAM15 (5 mg/kg i.p.) after the mice became ill (6 hours; at time of first dose of fluid and antibiotics) also increased survival (survival at 7 days, CLP+Vehicle: 15% vs CLP+BAM15: 44%, p<0.05) and improved kidney dysfunction at 18 hours after CLP (serum creatinine, CLP+Vehicle vs CLP+BAM15: 0.40 ± 0.08 vs 0.19 ± 0.06 mg/dl, p<0.05; BUN, CLP+Vehicle: 104.0 ± 14.4 mg/dl vs CLP+BAM15: 46.7 ± 8.2 mg/dl, p<0.05) (Figure 1D-F). BAM15 treatment did not have any apparent harmful effects on sham-treated, non-septic mice.

**Figure 1.**
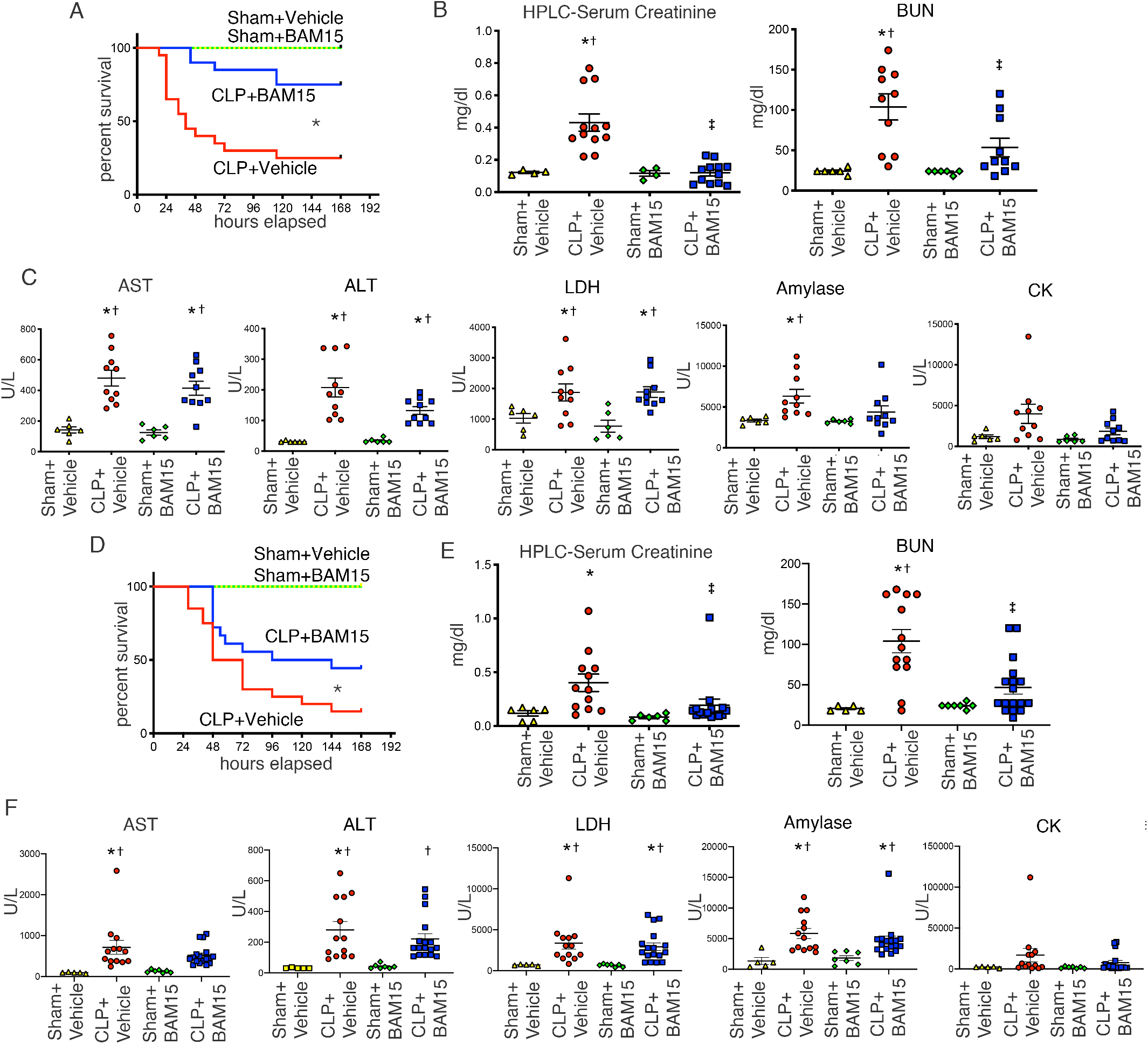
BAM15 treatment improves mortality and acute kidney injury (AKI) in septic mice, even with delay administration. (A) Kaplan-Meier curves of Sham or CLP male mice treated with vehicle (at 0 hours) and Sham or CLP treated with BAM15 (5mg/kg, at 0 hours) for 7 days. Sham+ Vehicle/ BAM15: n=4 each, CLP+ Vehicle/ BAM15: n=20 each. Log-rank test. *p<0.05, CLP + Vehicle vs other groups. (B-C) Serum creatinine measured by HPLC, BUN (B), AST, ALT, LDH, Amylase, and Creatinine Kinase (CK) (C) by biochemical examination at 18hrs after Sham or CLP treated with vehicle (at 0 hours) or BAM15 (5mg/kg, at 0 hours). All bars show mean ± SEM of each group (Sham + Vehicle: n=4~6, CLP + Vehicle: n=10~12, Sham + BAM15: n=4~6, CLP + BAM15: n=10~12). Sidak’s multiple comparisons test following one-way ANOVA test. *: vs Sham + Vehicle, p<0.05. †: vs Sham + BAM15, p<0.05. ‡: vs CLP + Vehicle, p<0.05. (D) Kaplan-Meier curves for 7 days of Sham or CLP male mice treated with vehicle (at 6 hours) or BAM15 (5mg/kg, at 6 hours). Sham+ Vehicle/ BAM15: n=4 each, CLP+ Vehicle/ BAM15: n=20, 19. Log-rank test. * CLP + Vehicle vs other groups, p<0.05. (E-F) Serum creatinine measured by HPLC and BUN (E) and AST, ALT, LDH, Amylase, and CK (F) by biochemical examination at 18 hours after Sham or CLP treated with vehicle (at 6 hours) or BAM15 (5mg/kg, at 6 hours). All bars show mean ± SEM of each group (Sham + Vehicle: n=5, CLP + Vehicle: n=13, Sham + BAM15: n=7, CLP + BAM15: n=17). Dunn’s multiple comparisons test following Kruskal-Wallis test. *: vs Sham + Vehicle, p<0.05. †: vs Sham + BAM15, p<0.05. ‡: vs CLP + Vehicle, p<0.05.

### BAM15 improved critical vital signs and reduced kidney hypoxia and damage in sepsis-AKI

We also monitored critical vital signs, including mean blood pressure (mBP), heart rate (HR), and body temperature (BT), in CLP mice and sham mice treated with vehicle or BAM15 (5mg/kg injected at 0 hours). These three physiological parameters decreased after CLP surgery in both vehicle and BAM15-treated mice compared with sham mice. However, BAM15 significantly blunted the reduction of mBP beginning 14 hours after injection, the reduction in HR beginning 9 hours after injection, and reduction in BT beginning 6 hours after injection. (Figure 2A).

**Figure 2.**
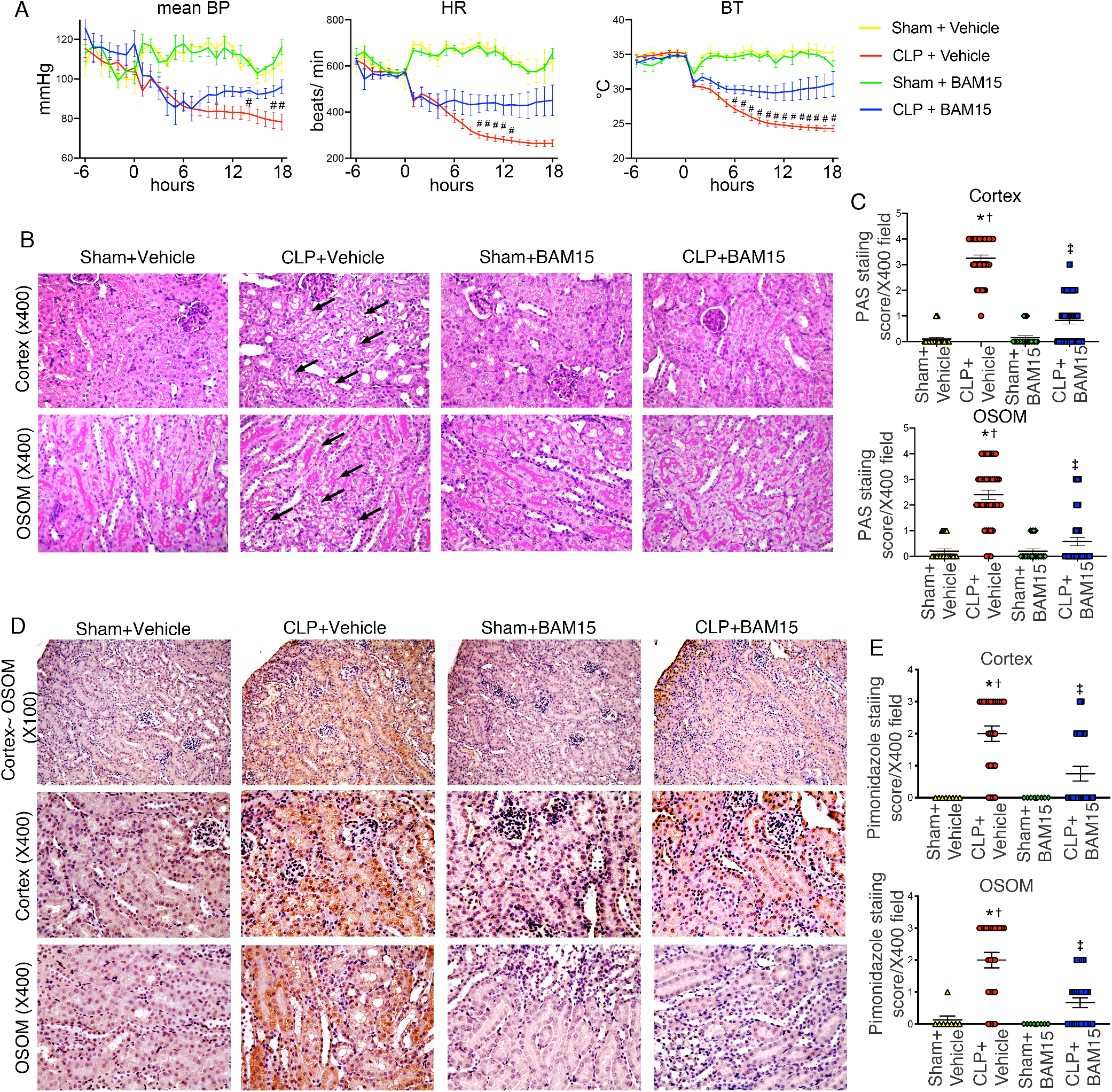
BAM15 improves septic shock and reduces injury and hypoxia in septic kidney. (A) Continuous mean blood pressure (mBP), heart rate (HR), and body temperature (BT) before and after Sham or CLP surgery with vehicle (at 0 hours) or BAM15 (5mg/kg, at 0 hours). Data were averaged across consecutive 1-hour windows. The bars show mean of ± SEM of 1-hour window-data of all mice in each group. CLP + Vehicle vs CLP + BAM15. Sham + Vehicle: n=3, CLP + Vehicle: n=9, Sham + BAM15: n=3, CLP + BAM15: n=8. Tukey’s multiple comparisons test following two-way ANOVA analysis. #: p<0.05, CLP+ Vehicle vs CLP + BAM15. (B-C) Periodic acid-Schiff staining of cortex and outer stripe of the outer medulla (OSOM) in kidney at 18 hours after Sham or CLP mice treated with vehicle (at 0 hours) or BAM15 (5mg/kg, at 0 hours). (B) Representative images of cortex and OSOM. Arrows show vacuolization in proximal tubule cells. Original magnification, x400. (C) Tubular damage score in cortex and OSOM of kidney at 18 hours after Sham (4 mice each, total 20 fields of x400) or CLP (8 mice each, total 40 fields of X400) treated with vehicle (at 0 hours) or BAM15 (5mg/kg, at 0 hours). All graph represent mean±SEM. Dunn’s multiple comparisons test following Kruskal Wallis test. *: vs Sham + Vehicle, p<0.05. †: vs Sham + BAM15, p<0.05. ‡: vs CLP + Vehicle, p<0.05. (D-E) Pimonidazole staining of cortex and outer stripe of the outer medulla (OSOM) in kidney at 18 hours after Sham or CLP mice treated with vehicle (at 0 hours) or BAM15 (5mg/kg, at 0 hours). (D) Representative images of cortex and OSOM. Original magnification, upper: X200, middle and lower: X400. (E) Tubular hypoxia score in cortex and OSOM of kidney at 18 hours after Sham or CLP treated with vehicle (at 0 hours) or BAM15 (5mg/kg, at 0 hours). Each group has 4 mice and total 24 fields for CLP groups and 8 fields for Sham groups. All graph represent mean±SEM. Dunn’s multiple comparisons test following Kruskal Wallis test. *: vs Sham + Vehicle, p<0.05. †: vs Sham + BAM15, p<0.05. ‡: vs CLP + Vehicle, p<0.05.

Kidney tissue damage from sepsis is detected histologically by cytoplasmic vacuoles in tubule cells (22). Cytoplasmic vacuoles were increased in proximal tubule cells of cortex and outer stripe of the outer medulla (OSOM) 18 hours after CLP, and was decreased by early treatment with BAM15 (5mg/kg i.p.). Injury scores were as follows; cortex, CLP+Vehicle: 3.3 ± 0.13 vs CLP+BAM15: 0.83 ± 0.14; OSOM, CLP+Vehicle: 2.4 ± 0.19 vs CLP+BAM15: 0.58 ± 0.16) (**Figure 2B-C**).

Tissue hypoxia is an important contributor to sepsis-induced organ dysfunction. Proximal tubule hypoxia was assessed by pimonidazole incorporation *in vivo* (24); pimonidazole reacts with cellular proteins at oxygen tensions < 10 mmHg. Pimonidazole incorporation was detected in proximal tubule cells in the cortex and the OSOM at 18 hours after CLP and was decreased by early treatment with BAM15 (5mg/kg i.p.) (hypoxia score; CLP+Vehicle: 2.0 ± 0.24 vs CLP+BAM15: 0.75 ± 0.23; OSOM, CLP+Vehicle: 2.0 ± 0.24 vs CLP+BAM15: 0.67 ± 0.16, p<0.05) (**Figure 2D-E**). Thus, BAM15 treatment reduced both sepsis-induced kidney tissue injury and tissue hypoxia.

### BAM15 reduced serum cytokines and immunosuppression

Sepsis causes both stimulation and suppression of the immune system. As expected, CLP increased serum IL-6, IL10, TNF-α and IL-17 (Figure 3A). BAM15 inhibited increases in both serum IL-6 and IL-10. Splenic apoptosis provides a measure of immunosuppression in mouse models of sepsis. CLP increased cleaved caspase-3 positive cells in spleen at 18 hours, which was decreased by pretreatment with BAM15 (5mg/kg i.p. at 0 hours after CLP) (positive cells, CLP + Vehicle: 26 ± 2.9/high power field (HPF) vs CLP + BAM15: 8.0 ± 1.6/HPF, p<0.05) (Figure 3B-C). Thus, BAM15 inhibited both the overproduction of systemic cytokines and immunosuppression after sepsis.

**Figure 3.**
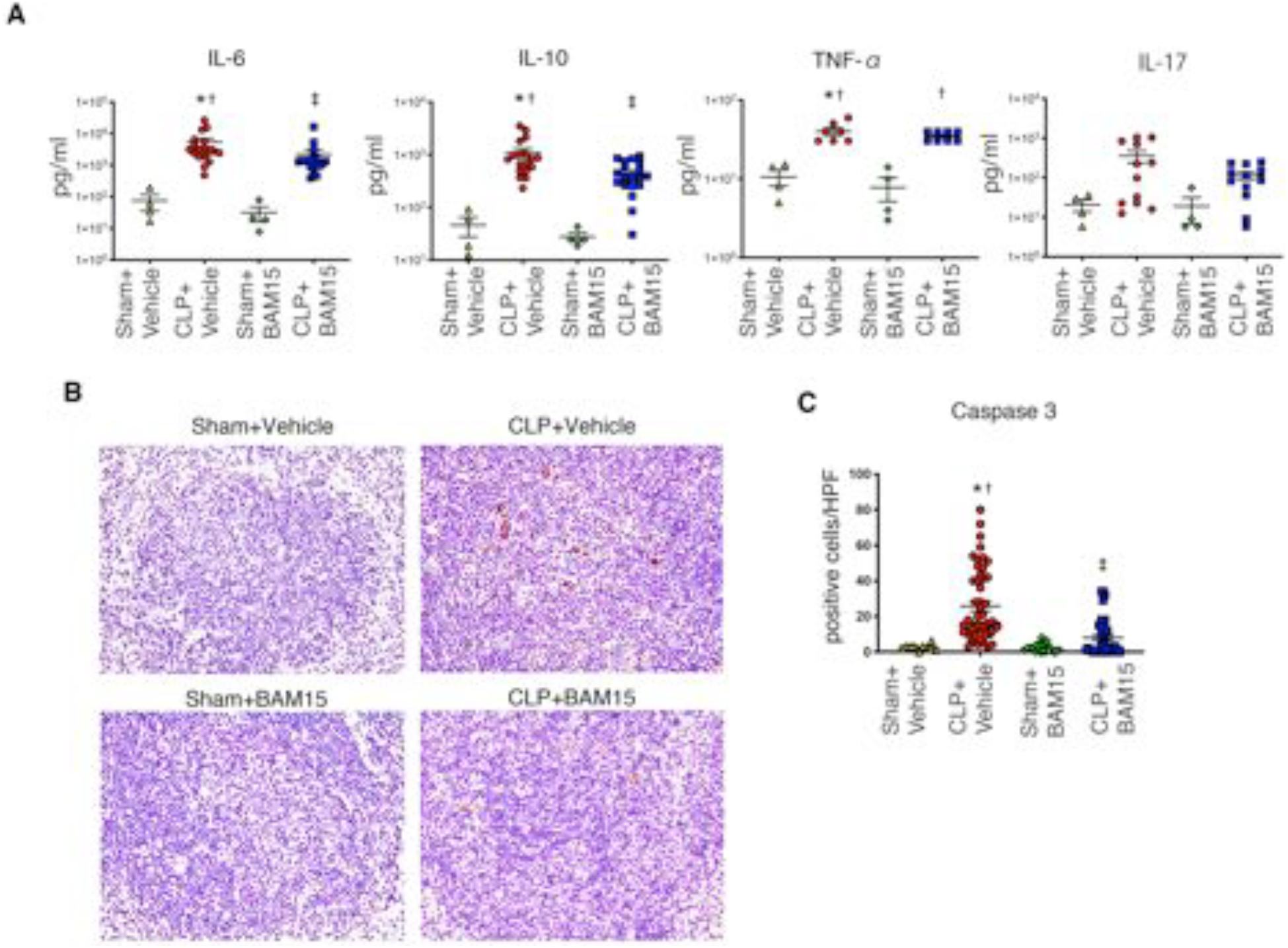
BAM15 inhibits splenic apoptosis and production of some cytokines. (A) Cleaved caspase 3 staining in spleen at 18 hours after Sham or CLP mice treated with vehicle (at 0 hours) or BAM15 (5mg/kg, at 0 hours). Original magnification, X400. (B) Positive cells of cleaved caspase 3 in spleen of Sham (4 mice each, total 20 fields of x400) or CLP (8 mice each, total 40~50 fields of X400) treated with vehicle (at 0 hours) or BAM15 (5mg/kg, at 0 hours) at 18 hours. (C) Cytokines in serum at 18 hours after Sham or CLP mice treated with vehicle (at 0 hours) or BAM15 (5mg/kg, at 0 hours). n=4, 7~18, 4, 8~19 mice. The bar of all graphs represent mean±SEM. Dunn’s multiple comparisons test following Kruskal Wallis test. * vs Sham + Vehicle, p<0.05. † vs Sham + BAM15, p<0.05. ‡ vs CLP + Vehicle, p<0.05.

### BAM15 is neither bacteriostatic nor bactericidal

Bacteria also have a membrane electron transport chain (25); thus, BAM15 might have a direct effect on bacterial energetics. We evaluated the effect of BAM15 on pathogens in the peritoneal cavity of septic mice. We found that BAM15 (5 mg/kg) given to septic mice at 0 hours after surgery did not alter the number of bacterial colonies in blood or abdominal fluid assessed 18 hours after CLP. Blood bacterial counts were as follows: CLP + Vehicle: 1.1×10^4^ ± 0.90×10^4^ CFU/ml vs CLP + BAM15: 0.10×10^4^ ± 0.053 × 10^4^ CFU/ml, p=0.4. Abdominal bacterial counts were as follows: CLP + Vehicle: 1.2×10^5^ ± 0.92×10^5^ CFU/ml vs CLP + BAM15: 6.9×10^5^ ± 6.6 × 10^5^ CFU/ml, p>0.99) (**Supplemental Figure 2A-B**). We also tested the bactericidal ability of BAM15 *in vitro* by directly adding BAM15 (0, 1, 5, 10, 20, and 50 μM) to a suspension of cecal material. BAM15 did not alter the number of bacterial colonies at any concentration (**Supplemental Figure 2C**). We conclude that BAM15 does not have bactericidal activity *in vivo* or *in vitro*.

### BAM15 inhibited cell-free mtDNA in mice

While cfmtDNA is a biomarker of tissue injury in critically ill patients (26), it is unknown whether cfmtDNA levels can be used to predict and/or monitor drug efficacy. We evaluated the drug-biomarker relationship between BAM15 and cfmtDNA in male mice. Plasma cfmtDNA was increased from 2 to 18 hours after CLP, and that BAM15, given at the time of CLP surgery, reduced cfmtDNA 2 to 18 hours after CLP (plasma cfmtDNA at 2 hours, CLP + Vehicle: 1.4×10^5^ ± 0.26×10^5^ copies/μl vs CLP + BAM15: 0.32×10^5^ ± 0.09 × 10^5^ copies/μl, p<0.05) (**Figure 4A**). Urinary cfmtDNA was also increased beginning 6 hours after CLP, and BAM15, given at the time of CLP surgery, reduced urinary cfmtDNA after CLP (mtDNA/gCr in urine at 6 hours, CLP + Vehicle: 1.8×10^6^ ± 1.0×10^6^ copies/μl vs CLP + BAM15: 0.52×10^5^ ± 0.14 × 10^5^ copies/μl, p<0.05) (**Figure 4B-C**).

**Figure 4.**
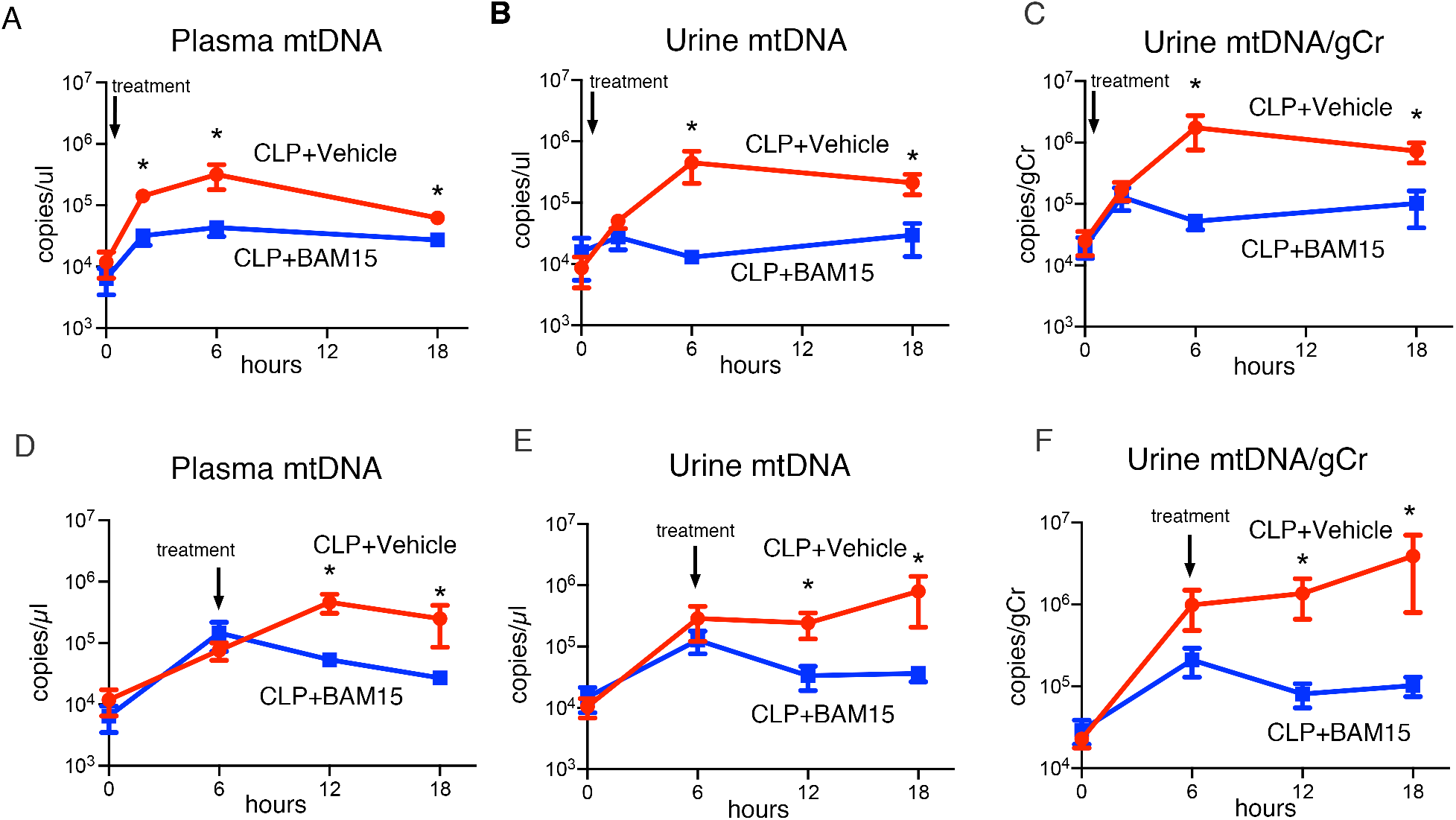
Both early and delay BAM15 treatment decreases circulating mtDNA. (A-C) Time course of plasma mtDNA level (A) and urine mtDNA (B) and urine mtDNA adjusted to the creatinine excretion (C) at 18 hours after CLP (n=8 each) mice treated with vehicle (at 0 hours) or BAM15 (5mg/kg, at 0 hours). (D-F) Time course of plasma mtDNA level (D) and urine mtDNA (E) and urine mtDNA adjusted to the creatinine excretion (F) in CLP mice (n=8 each) treated with vehicle (at 6 hours) or BAM15 (5mg/kg, at 6 hours). All graph represent mean ± SEM. Analysis between groups at each time point was performed with Sidak’s multiple comparisons test following mixed-effects analysis. *: p<0.05.

The effect of BAM15 treatment on cfmtDNA was also observed in female mice (**Supplemental Figure 1D-F**). Delayed treatment with BAM15 (6-hour delay) also inhibited both plasma and urine cfmtDNA (plasma cfmtDNA at 12 hours, CLP + Vehicle: 4.6×10^5^ ± 1.6×10^5^ copies/μl vs CLP + BAM15: 0.53×10^5^ ± 0.12 × 10^5^ copies/μl, p<0.05; urinary mtDNA/gCr at 12 hours, CLP + Vehicle: 1.4×10^6^ ± 0.70×10^6^ copies/gCr vs CLP + BAM15: 0.81×10^5^ ± 0.26 × 10^5^ copies/gCr, p<0.05) (**Figure 4D-F**). These results suggest that plasma and urinary cfmtDNA levels are responsive to BAM15 treatment in sepsis-AKI; thus, BAM15 and cfmtDNA are a potential drug-companion biomarker pair.

### BAM15 accelerates mitochondrial respiration without ATP in cultured renal tubule cells

We investigated the direct impact of BAM15 on kidney tubule cell mitochondria (*i.e.*, in the absence of systemic mediators). First, we determined the optimal concentration of BAM15 as a mitochondrial uncoupler in kidney tubule cells by measuring oxygen consumption rate (OCR) and mitochondrial respiratory function in primary cultured proximal tubule cells from healthy CD-1 mice. BAM15 (1, 2, 5, 10, 20, 50 and 100 μM concentrations) functioned as a mitochondrial uncoupler similar to FCCP (the most commonly used uncoupler). There was a concentration-dependent increase in maximal respiration following addition of 1 μM oligomycin, an ATP synthase inhibitor (Supplemental Figure 3A-D). Maximal respiration peaked at 20 μM BAM15, which matched the level in response to 5 μM FCCP. 20 μM BAM15 increased maximal respiration more than 20 μM FCCP (p<0.05) (Supplemental Figure 3E).

### BAM15 inhibits overproduction of mitochondrial superoxide in cultured renal tubule cells exposed to serum from septic mice

We established an *in vitro* model of septic kidney tubule cells by incubating primary cultured tubule cells with serum from septic mice obtained 18 hours after CLP. Overproduction of mtROS has been proposed as a pathophysiologic mechanism for sepsis-AKI (27). Mitochondrial uncouplers paradoxically decrease mitochondrial superoxide production by increasing the electron transfer rate, which decreases the dwell time for single electrons traversing the electron transport chain (ETC) (15–17).

To evaluate the effect of BAM15 on renal mtROS, we measured the production of mitochondrial superoxide using mitoSOX-red in primary cultured proximal tubule cells co-incubated with serum from CLP mice (‘septic tubule cells’); measurements were made at 0, 6, 12, and 24 hours following the addition of mouse serum. Co-incubation with septic serum increased mitoSOX-Red intensity over time, compared to co-incubation with control serum. BAM15 (either 10μM or 20μM) inhibited the overproduction of superoxide induced by CLP serum (**Figure 5A and B**), with a significantly larger effect at 20μM BAM15. These results suggest that BAM15 can act, in part, by decreasing mtROS production in tubule cells during sepsis.

**Figure 5.**
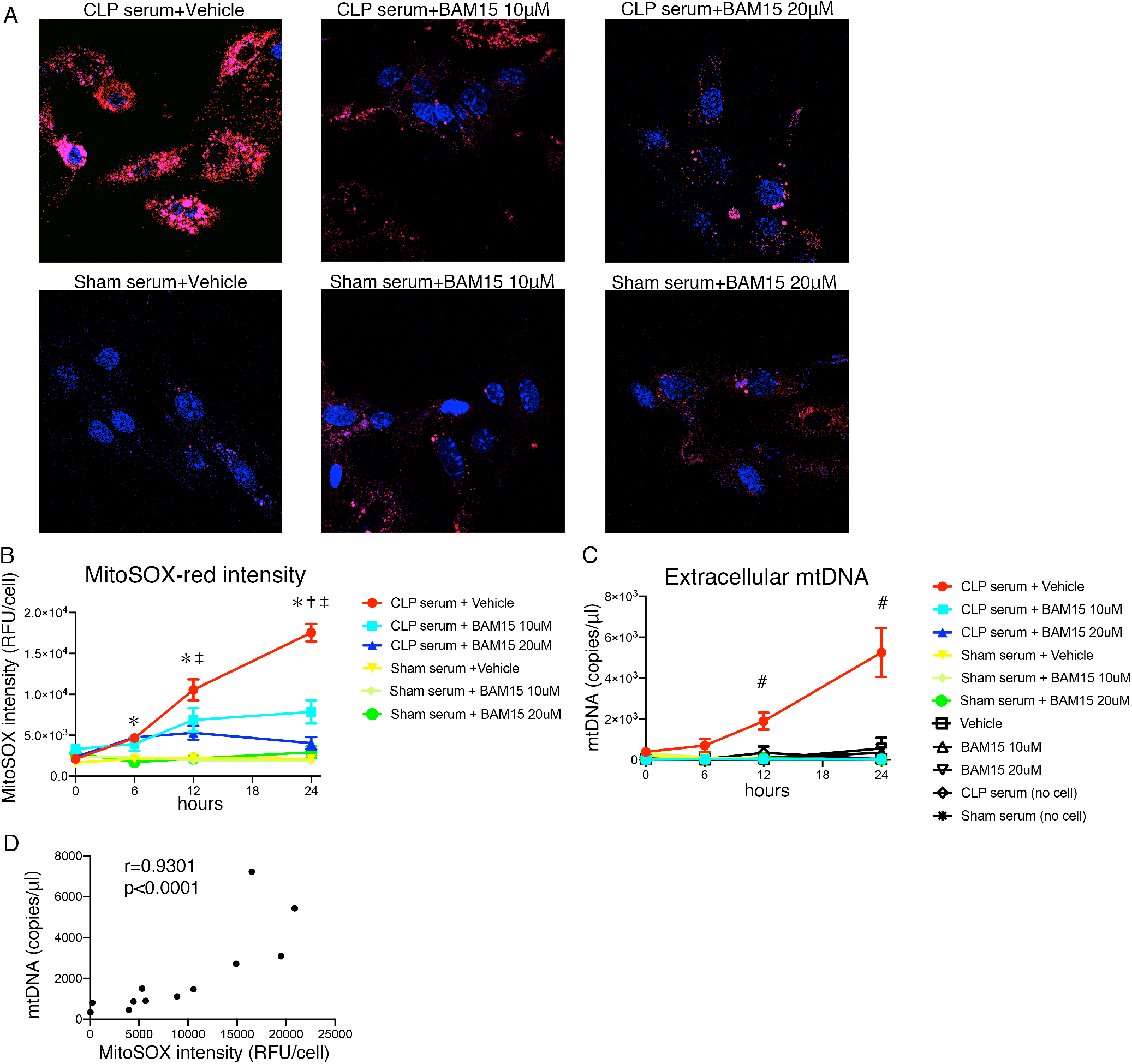
BAM15 inhibits production of mitochondrial reactive oxygen species (ROS) in proximal tubular cells (PTCs) linked with decreasing mtDNA released from PTCs. (A-B) Serial live cell imaging of mitochondrial ROS in mouse primary PTCs incubated with Sham or CLP serum treated with vehicle or BAM15 (10μM or 20μM). (A) Representative images of mitochondrial ROS of primary PTCs in each group at 24 hours after incubation. Red: MitoSOX-red, Blue: Hoechst 33342. Original magnification, x400. (B) Time course of fluorescence intensity of MitoSOX-red serially measured. All graph represent mean±SEM. Tukey’s multiple comparisons test following two-way ANOVA test. N=18~31, 16~24, 19~24, 18~32, 9~13, 8~13 on 3 biological replicates per each condition. *: vs each Sham serum group, p<0.05. †: vs CLP serum + 10μM BAM15, p<0.05. ‡: vs CLP serum + 20μM BAM15, p<0.05. (C) Time course of extracellular mtDNA level of the supernatant of (B). Controls were medium with CLP/Sham serum or supernatants of PTCs treated with only Vehicle/BAM15. #: CLP serum + Vehicle vs each and every other group, p<0.05. Tukey’s multiple comparisons test following two-way ANOVA test. (D) Correlation between extracellular mtDNA level and the corresponding mitoSOX-red intensity in the primary PTCs incubated with CLP serum treated with vehicle. n=8 from 0, 6,12 and 24h on 2 biological replicates. r value is Pearson’s correlation coefficient, *p<0.05.

### BAM15 inhibits mtDNA release from septic tubule cells

To evaluate whether mtDNA is released from septic tubule cells, we measured the appearance of cell-free mtDNA into the media of cultured tubule cells at 0, 6, 12, and 24 hours after incubation with septic serum. The concentration of cfmtDNA increased over time. Both 10 and 20 μM BAM15 inhibited cfmtDNA release from septic tubule cells. (Figure 5C). Furthermore, cfmtDNA release correlated with MitoSOX generation in septic tubule cells (r=0.93, p<0.0001) (Figure 5D). The purified mtDNA from livers from mice subjected to CLP caused an increase in mtROS of tubule cells (Supplemental Figure 4), These results suggest that the BAM15 inhibition of superoxide production and mtDNA release from tubule cells are linked mechanistically.

### BAM15 inhibits overproduction of reactive nitrogen species in septic mouse kidneys

We measured kidney reactive nitrogen species (RNS) in mice subjected to CLP with BAM15 to determine the effect of BAM15 on renal oxidants *in vivo*. Superoxide (O_2_^-^) can react with nitric oxide (NO) to form peroxynitrite (ONOO^-^). Peroxynitrite is a powerful cytotoxic oxidant in septic kidneys (28) that can oxidize thiols and DNA bases, or can modify proteins and lipids by nitration. Immunostaining for nitrotyrosine, a product of tyrosine nitration, was performed on kidney sections from mice subjected to CLP and BAM15 administration (at 0 hours) at each timepoint (2, 6, 18 hours after CLP). Nitrotyrosine was found in the tubular epithelial cells from 2 hours (cortex) and 6 hours (OSOM) after mice subjected to CLP and administered with vehicle. However, BAM15 treatment decreased nitrotyrosine in tubule cells of cortex and OSOM (Figure 6). Our result confirmed that BAM15 decreased cytotoxic oxidants in septic kidney *in vivo*, in addition to decreasing superoxide levels in tubules.

**Figure 6.**
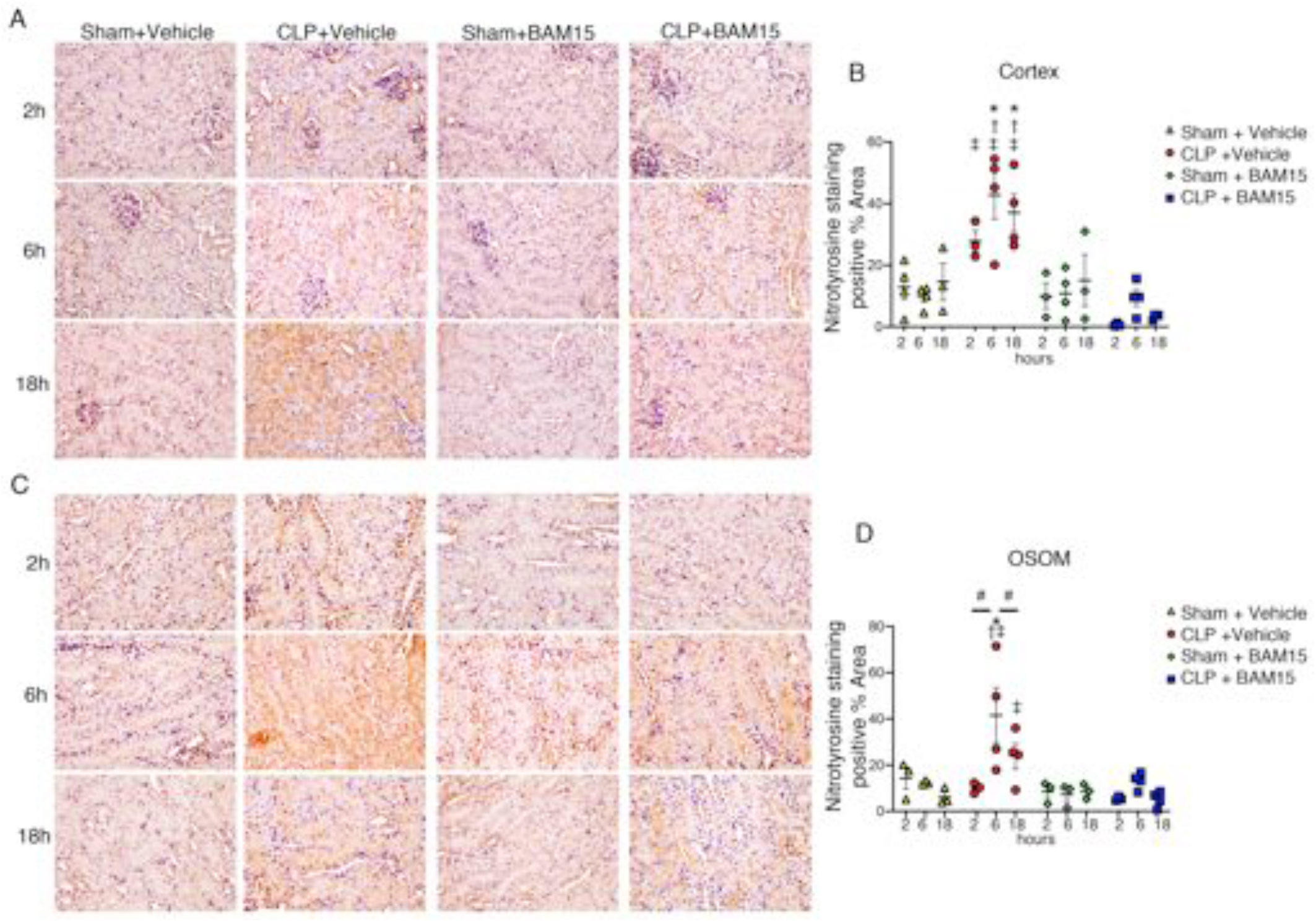
BAM15 inhibits production of reactive nitrogen species (RNS) in PTCs of septic kidney. (A-D) Representative Nitrotyrosine images and the positive area rate of cortex (A, B) and OSOM (C, D) in kidney at 2, 6 and 18 hours after Sham or CLP treated with vehicle (at 0 hours) or BAM15 (5mg/kg, at 0 hours). Original magnification, x400. The positive area rate was calculated by Fiji/ Image J software. Each dot represents the average of positive area of 3-4 fields of the mouse kidney (3-4 mice/ group). The bar shows mean ± SEM. Tukey’s multiple comparisons test following two-way ANOVA test. *: vs Sham + Vehicle, p<0.05. †: vs Sham + BAM15, p<0.05. ‡: vs CLP + Vehicle, p<0.05. #: comparison between time points, p<0.05.

### BAM15 effects on mitochondrial biogenesis in septic kidney

We also evaluated the effect of BAM15 on mitochondrial biogenesis after CLP. PGC1α is a transcription factor that regulates expression of multiple genes involved in mitochondrial biogenesis and is highly expressed in healthy renal tubules (20). Kidney PGC1α protein expression decreased within 2 hours after mice were subjected to CLP and vehicle administration, compared with mice subjected to sham surgery and vehicle administration, and PGC1α protein levels did not return to baseline. However, administration of BAM15 prior to CLP increased the expression of PGC1α at 6 hours after CLP, compared with mice subjected to CLP and administered with vehicle, and PGC1α levels approached baseline values over time (Figure 7A).

**Figure 7.**
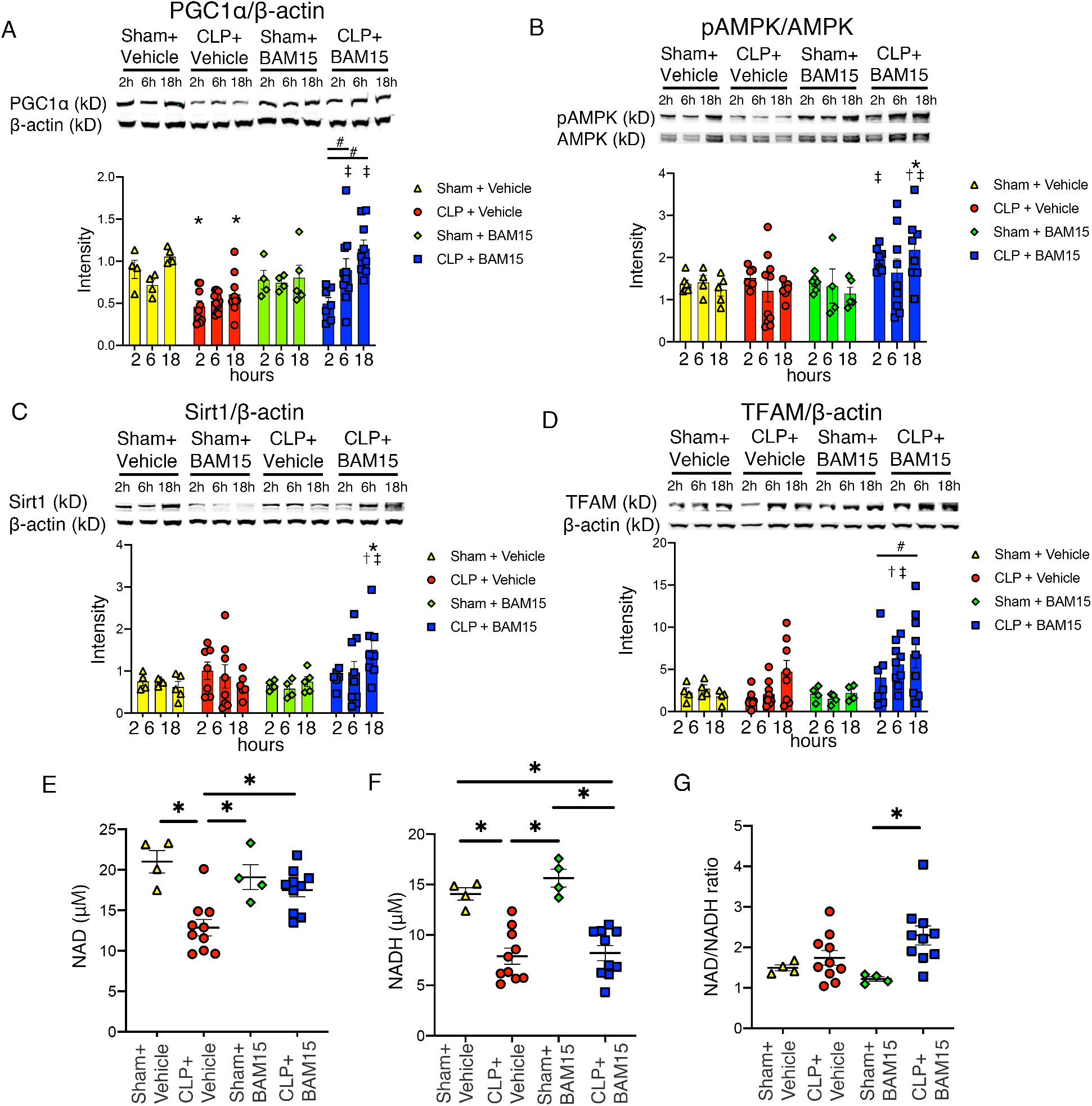
BAM15 recovers mitochondrial biogenesis in septic kidney via PGC1-α with increasing its regulator, AMPK and Sirt1 and increasing TFAM. (A-D) Representative blot image and fluorescent intensity of western-blotting of phosphorylated AMPK/AMPK and AMPK (A), Sirt1 (B), PGC1-α(C) and TFAM (D) in whole kidney at 2, 6 and 18 hours after Sham or CLP mice treated with vehicle (at 0 hours) or BAM15 (5mg/kg, at 0hrs). β-actin is internal control of (B-D). n=4-11 mice per each group. All bar represent mean ± SEM. Tukey’s multiple comparisons test following two-way ANOVA test. *: vs Sham + Vehicle, p<0.05. †: vs Sham + BAM15, p<0.05. ‡: vs CLP + Vehicle, p<0.05. #: comparison between time points, p<0.05. (E-G) Concentration of NAD^+^ (E), NADH (F), NAD^+^/NADH ratio (G) in cortex of kidney at 18 hours after Sham or CLP mice treated with vehicle (at 0 hours) or BAM15 (5mg/kg, at 0 hours). n=4 for each sham group, n=10 for each CLP group. All graph represent mean ± SEM. Tukey’s multiple comparisons test following one-way ANOVA test. *: p<0.05.

PGC1α is activated by AMP-activated protein kinase (AMPK) and sirtuin 1 (SIRT1) (29, 30). We found that CLP depressed PGC1α without significant changes in AMPK and SIRT1 levels; pre-treatment with BAM15 increased expression of phosphorylated AMPK at 2 to 18 hours and SIRT1 at 18 hours (**Figure 7B-C**). Thus, BAM15 appears to activate AMPK during the early phase of sepsis-AKI and activates SIRT1 during the late phase of sepsis-AKI, followed by increasing mitochondrial biogenesis associated with PGC1α activation.

We also measured NAD^+^ in kidney at 18 hours after CLP. The activation of PGC1α by SIRT1 requires NAD^+^ (31). Levels of NAD^+^, as well as NADH, were decreased in mice subjected to CLP and administered with vehicle, which did not change the NAD^+^/NADH ratio (**Figure 7E-G**). Early BAM15 pre-treatment partially restored NAD^+^ (**Figure 7E**), but did not alter NADH (**Figure 7F**). Our data support an activation of Sirt1 by BAM15 through increased NAD^+^ levels, which would be expected to restore PGC1α levels.

PGC-1a promotes expression of many nuclear genes whose products are imported into mitochondria, including the mitochondrial transcription factor A (TFAM). TFAM is a mtDNA-binding protein that is essential for genome maintenance (32) and plays a central role in the mtDNA stress-mediated inflammatory response, including to AKI (33). BAM15 treatment increased TFAM from 6 hours after CLP (**Figure 7D**) compared with CLP treated with vehicle. These results are consistent with BAM15 treatment enhancing expression of these mitochondrial biogenesis-related proteins.

## Discussion

The pathogenesis of multi-organ failure in sepsis is complex and the underlying pathways are heterogeneous. Multiple PAMPs and DAMPs released from injured tissues can cause a runaway inflammatory response, which propagates additional organ damage and can be immunosuppressive (34). Many PAMPs and DAMPs are released during sepsis in animals and humans (35), suggesting that individual targeted approaches might be ineffective, or that therapy should target a common upstream node. We previously showed that mice lacking Toll-like receptor 9 (TLR9) or treated with a TLR9 inhibitor were protected from sepsis (8, 36) and that a TLR9 ligand cfmtDNA is released during early sepsis and can contributes to cytokine production and AKI (8). Inhibition with a TLR9 inhibitor improved survival (36). It is not clear whether this pathologic pathway is amenable to drug targeting, as TLR9 inhibition did not alter plasma mtDNA levels (8). Moreover, there are other sensors for cfmtDNA besides TLR9 (37). We hypothesized that the optimal companion biomarker pairs for sepsis and sepsis AKI would be cfmtDNA and BAM15, a mitochondrial protectant. Here, we show that 1) BAM15 improved mortality and AKI in sepsis even with delayed treatment, 2) levels of one DAMP, cfmtDNA, are diminished by BAM15 treatment *in vivo* and *in vitro*, 3) cfmtDNA functions as an early drug efficacy biomarker, and 4) mtDNA causes mitochondrial and tissue injury via production of superoxide.

In this study, the mitochondrial uncoupler BAM15 improved survival when given at the time of CLP surgery. Survival also improved when treatment was delayed 6 hours, well after the animals exhibited clinical signs of sepsis (**Figure 1**). Efficacy with delayed treatment is critical for drug development, as the early stages of human sepsis may begin out of hospital and are often difficult to diagnose. Effective therapeutics are needed for overt sepsis, including septic shock. BAM15 joins a short list of therapeutic agents that improve survival, rather than just delaying death, in animal models of sepsis (38–41). In these studies, the actions of BAM15 were synergistic with those of fluids and antibiotics, which are typically given to septic patients. Unlike other uncouplers that are bactericidal (42), BAM15 did not have antibacterial effects, as assessed by bacterial counts in blood and peritoneal fluid (**Supplemental Figure 2**). This observation is consistent with a lack of direct impact on bacterial growth *in vitro* and *in vivo*. Thus, BAM15 acts on one or more critical early upstream events or only a subset of mediators of sepsis pathophysiology.

BAM15 had both systemic and renal effects. BAM15 improved systemic hemodynamics (reducing hypotension, increasing HR, and BT), and decreased splenic apoptosis (a marker of immunosuppression) and some systemic cytokine levels (IL-6, IL-10). Interestingly, BAM15 did not have much effect on other organ injury nor did it alter TNFα or IL-17 levels. The lack of effect on TNFα is puzzling, as a previous report found that BAM15 reduced increased expression of TNF-α in retinal tissues during donor organ transportation (43).

BAM15 protected against AKI, assessed by measures of glomerular filtration function (serum BUN and creatinine), histologic damage indicative of sepsis (tubular vacuolization, tubule dilation) and tubule hypoxia (pimonidazole staining, **Figure 2D-E**). The histologic protection conferred by BAM15 differs from its effect on renal ischemia/reperfusion injury, where it prevented tubular necrosis and vascular pooling (13). The major findings reported here were similar in male and female mice, indicating that the benefit from BAM15 treatment is not sex-specific. BAM15 protected the kidney from AKI and reduced mortality, even when given during established sepsis. Surprisingly, BAM15 did not manifest widespread protection against other organ dysfunction and only affected some circulating mediators of sepsis pathophysiology.

We and others have investigated many potential therapeutic agents for sepsis. However, the sepsis field has lacked a precision medicine approach to selecting the right drug, timing, and dose for an individual patient. Our previous work on the TLR9 (8, 36) and mtDNA (8) pathway had determined that circulating mtDNA is released during an early phase of sepsis, induces cytokine production and splenic apoptosis, and causes renal tubular injury (8). Others have also found mitochondrial damage in sepsis and sepsis AKI (20, 29, 44–46). Infusion of damaged mitochondria produce a sepsis-like kidney damage by histology that is reversed by DNase, suggesting mtDNA is toxic (8). In a study of patients with COVID-19, we found that human mtDNA caused proximal tubule damage *in vitro* (11). In the current study, we showed that circulating mtDNA and urinary mtDNA are increased very early in sepsis. In mouse studies, urinary mtDNA was more sensitive than BUN in detecting subclinical renal injury in an ischemia/reperfusion model (47). Small clinical studies show that high levels of circulating mtDNA, and especially urinary mtDNA, predict AKI and mortality in critically ill patients (48) and are associated with more severe sepsis-AKI (46).

Importantly, we showed that BAM15 decreased circulating and urinary cfmtDNA after administration of BAM15 both *in vivo* in male and female mice (**Figure 4, Supplementary Figure 1**), and in an *in vitro* model using septic serum and mouse kidney tubules as detector cells (**Figure 5**). As mtDNA levels changed rapidly after BAM15 administration, changes in mtDNA levels might be able to predict drug efficacy more quickly than other approaches, such a serum creatinine or cytokine levels. By contrast, TLR9 deficiency during sepsis, in TLR9 knock-out mice, altered peritoneal fluid mtDNA levels, but not plasma mtDNA levels, compared to wild-type mice (8). Whereas TLR9 inhibition and plasma mtDNA do not function as a drug-companion biomarker pair during sepsis, BAM15 and plasma (and urine) mtDNA do function in this way.

The effects of sepsis on mtDNA, and mtDNA on tissue damage were reproduced in an *in vitro* septic kidney tubule assay. Serum from septic mice increased mtROS and also increased mtDNA concentration in culture medium, and both were reduced by BAM15 (**Figure 5**). The amount of extracellular mtDNA detected from 6 to 24 hours was substantially higher than mtDNA induced by exposure to septic serum. The tight correlation between mtDNA concentration and fluorescence intensity of mitoSOX Red suggests a possible mechanistic relationship between ROS overproduction and mtDNA release. In this model, ROS cause mtDNA release, and mtDNA would cause additional ROS production, thus forming a positive feedback loop. However, we have not determined the kinetics of ROS production leading to mtDNA release, but mtDNA purified from liver of mice subjected to CLP can cause an increase in mtROS (**Supplemental Figure 4**), as we showed for cell free DNA (cfnDNA) including cfmtDNA purified from COVID-19 patient plasma(11). Even so, the indirect (mtDNA) and direct (mtROS) toxic mediators apparently work in tandem. The association could be caused by release of other circulating PAMPs and DAMPs induced during sepsis, including perhaps genomic DNA. Previously, we showed that in COVID-19 patients, both cell-free mitochondrial and nuclear DNA (cfnDNA) was increased 10-100 fold in plasma, and the cfDNA came from multiple cell types including kidney (11). Circulating cfnDNA is also increased in septic patients (49). We have not evaluated the precise mechanism of the biomarker-drug relationship between nuclear DNA and BAM15 in septic mice; however, mtDNA and BAM15 would be expected to have a more intimate proximity relationship because BAM15 is a mitochondrial protectant. Taken together, our data support BAM15-mtDNA being tested in the future as a drug-biomarker pair to determine clinically which patients to treat, what dose to administer, how to adjust the dose, and when to terminate administration.

Sepsis is associated with mitochondrial DNA damage and reduced mitochondrial mass in the kidney of patients with sepsis-AKI (50). BAM15 is a mitochondrial uncoupler and postulated to protect mitochondria from oxidative stress induced damage during renal ischemia/reperfusion (13). However, the actual mechanism of BAM15 effect on kidney had not been determined. We found that BAM15 decreased reactive nitrogen species in septic kidneys (**Figure 6**), decreased mitochondrial superoxide production in septic renal cells (**Figure 5**), and normalized pathways that would inhibit mitochondrial biogenesis (**Figure 7**). BAM15 inhibited reactive nitrogen species in septic kidneys and superoxide overgeneration in septic tubule cells. Single electrons in the electron transport chain exist only transiently at the redox centers, and the dwell time for single electrons in an unstable state increases the likelihood of superoxide production (17). Mitochondrial uncouplers decrease mitochondrial superoxide production by stimulating faster electron transfer through the electron transfer chain. We showed superoxide generation and mtDNA release was correlated in septic renal tubule cells. Released mtDNA could act as DAMPs via TLR9 in the cytosol or extracellularly (7, 8), causing cell stress, including mtROS. Both mtROS and mtDNA might be involved as mediators in a dual positive feedback loop (described above), while other existing biomarkers of AKI (e.g., KIM-1, L-FABP) are associated with excessive ROS and hypoxia, respectively, are acute markers of tissue injury (51, 52), but are not known to mediate further damage.

Beyond inhibition of mtROS overproduction, BAM15 also altered several mediators of mitochondrial biogenesis, including PGC-1α (early and late post-CLP), pAMPK (early), SIRT1 (late), and NAD+ (late) (**Figure 7**). PGC-1α is a key transcriptional regulator of mitochondrial function, biogenesis, and respiration in many tissues, including kidney (53). Activated pAMPK and Sirt-1 can activate PGC1α in a dependent or independent manner with each other (54). The activation of PGC1α by SIRT1 requires NAD^+^-dependent deacetylation, and PGC1α, in turn, promotes NAD^+^ biosynthesis via the de novo biosynthetic pathway (31, 55). A previous study showed that lipopolysaccharide (LPS) suppresses the expression of renal PGC-1α, and that the decrease in PGC-1α correlates with the degree of renal impairment (55). Administration of an AMPK (56) or Sirt1 (57) activator was reported to improve multiple organ failure or mortality in sepsis rodent model. In our study, BAM15 treatment, as reported in other cell types (18, 58), activated AMPK in the kidney at an early phase of sepsis (31, 55). Moreover, BAM15 increased the expression of kidney Sirt1 during a late phase associated with an increase in renal NAD^+^. These results suggest that BAM15 may activate AMPK at early phase of sepsis-AKI and increase mitochondrial biogenesis by increasing PGC1α. The increased PGC1α might increase NAD^+^ which then activates Sirt1, and this restores PGC1α levels during the late phase of sepsis-AKI.

We also showed BAM15 increased TFAM expression in kidney during an early phase of sepsis. Sepsis is associated with mtDNA damage and a reduced mitochondrial mass in the kidney of patients with sepsis-AKI (50). PGC-1a activates expression of nuclear respiratory factor (NRF)-1 and −2, which together increase transcription of many genes whose products are imported into mitochondria, including TFAM (32, 59). The maintenance of mtDNA stability is primarily mediated by TFAM, an essential mtDNA packaging protein that is required for mtDNA replication and transcription (32). Aberrant mtDNA packaging induced by TFAM depletion may promote leakage of mtDNA into the cytosol, followed by activation of innate immune response (60), perhaps via TLR9 (36, 61). Indeed, Zhao et al. recently showed that mtROS could promote IRI-induced AKI by suppressing TFAM transcription and enhancing protease-mediated TFAM degradation (44). Thus, decreased renal mtROS by BAM15 may promote TFAM expression in the septic kidney and inhibit sepsis-AKI progression, perhaps via release of mtDNA.

There are limitations to the present study. We tested only one dose (5mg/kg i.p.) of BAM15, a dose suggested in the published literature (13). To investigate the optimal dose to other tissues, further pharmacokinetic studies are warranted. Further, it is unclear why BAM15 improved survival without much effect on organ dysfunction other than AKI, or effects on TNFα.

In conclusion, BAM15 administration either before or after induction of sepsis decreased mortality in a clinically-relevant animal model of abdominal sepsis that manifests bacteremia, unstable hemodynamics and a high mortality, despite treatment with volume resuscitation and broad-spectrum antibiotics. Even though BAM15 did not have bactericidal effect, BAM15 improved systemic hemodynamics, kidney injury, and attenuated splenic apoptosis, although it did not have effects on other organs or some cytokines. More critically, we established a tight coupling between the BAM15 effect on mtROS and mtDNA using *in vitro* assays, and effects on mitochondrial homeostasis. We conclude that BAM15 may be an effective preventive and therapeutic agent in sepsis, in part, by altering the trajectory of mitochondrial homeostasis. Our results suggest that BAM15 and mtDNA are mechanistically linked via mtROS, and may form a drug-companion biomarker pair as part of an evolving precision medicine approach to treating sepsis.

## Methods

### Animals

We followed National Institutes of Health (NIH) guidelines for the use and treatment of laboratory animals, and the NIDDK Animal Care and Use Committee approved all procedures. All animals had free access to water and chow throughout the study. We followed the recent recommendations on minimum quality threshold in pre-clinical sepsis studies (MQTiPSS) (62). Male or female CD-1 mice (Charles River Laboratories, Wilmington, MA) were 9-10 weeks old at the time of cecal ligation and puncture (CLP) surgery. CLP surgery was performed to induce sepsis, largely as previously described (22). After a midline incision, the cecum was located and ligated with 4-0 silk suture 15 mm from the tip, and a 21-g needle was passed through the ligated cecum. The cecum was returned to the peritoneal cavity and the incision closed. During surgery, a slow-release formulation of buprenorphine (0.5 mg/kg s.c.; SR Veterinary Technologies, Windsor, CO) was administered for analgesia (22). Mice were treated with vehicle (40 ml/kg of 3% DMSO diluted with 0.9% saline) or BAM15 (5mg/kg diluted with vehicle) i.p. at the time of surgery (0 hours) and then administered 40 ml/kg of 0.6% saline s.c. at 6 hours after CLP. In some mice, BAM15 treatment was delayed until 6 hours; these mice were given 40ml/kg 0.9% saline i.p. at the time of surgery and then administered 20ml/kg of vehicle or BAM15 (5mg/kg) i.p. with 20ml/kg of 0.6% saline s.c. at 6 hours after CLP. Antibiotics [14 mg/kg Primaxin (Imipenem and cilastatin), Merck, Whitehouse Station, NJ)] were given with fluids at 6 hours. In sham controls, surgery was performed as for CLP, with the cecum returned to the peritoneal cavity without ligation or puncture.

### Survival study

Mouse survival was assessed every 6 –12 hours after surgery. Antibiotic injection and fluid resuscitation were started at 6 hours after surgery by subcutaneous injection, and then repeated every 12 hours. Animals exceeding a threshold of morbidity were euthanized (63).

### Tubule cell culture

Renal tubular epithelial cells (TECs) were prepared from renal cell suspension of CD-1 mice following the method reported previously (64). In brief, kidneys were harvested, decapsulated, cut into small pieces with sterile instruments and digested in 30 mg/g tissue of collagenase type I (Millipore Sigma, Burlington, MA) for 30min at 37°C. The material was pushed through a sieve of 70 μm pore (BD, Franklin Lakes, NJ), washed and diluted in 2 mL of phosphate-buffered saline (PBS). The tubule segments were separated through 31% Percoll (GE Healthcare, Uppsala, Sweden) centrifugation at 800g for 10 min at 4°C. The pellet was collected and washed with PBS twice at 370 x g for 5 minutes at 4°C. Isolated TECs were cultured under sterile conditions at 37°C and 5% CO_2_ in conditioned Renal Epithelial Cell Growth Medium 2 (PromoCell, Heidelberg, Germany). The TECs isolated by this method have been reported to be mainly proximal tubule epithelial cells (65).

### Immunohistochemistry

After collection, tissue was immediately transferred to 10% formalin and fixed for 24 hours before paraffin embedding. Sections (4 μm) were stained with periodic acid-Schiff reagent. Semiquantitative assessment of kidney injury was performed by scoring 20 ~ 40 randomly selected, 40 objective fields per mouse on the following scale: 1) tubular damage observed in 0 –25% of the field, 2) 25–50%, 3) 50 –75%, and 4) 75–100%. For nitrotyrosine immunohistochemistry, sections were incubated overnight at 4°C in 1:100 dilution of rabbit polyclonal anti-nitro tyrosine antibody (ab42789; Abcam, Cambridge, MA). Slides were rinsed in PBS and developed using polyclonal goat anti-rabbit Immunoglobulins conjugated with horseradish peroxidase (HRP, Agilent Dako, Santa Clara, CA) as a secondary antibody, the manufacturer’s directions.

Splenic apoptosis was assessed by activated caspase 3 as described previously (61). Immunohistochemical staining was performed with anti- caspase 3 antibody (Cell Signaling Technology, Danvers, MA) as a primary antibody and polyclonal goat anti-rabbit Immunoglobulins conjugated with HRP (Agilent Dako, Santa Clara, CA) as a secondary antibody. The percent positive area in each 400x-magnified field was calculated using Fiji/ ImageJ software (National Institutes of Health, Bethesda, MD).

### Telemetry implantation and catheter insertion

For measuring blood pressure, heart rate, and body temperature measurements, mice were implanted with a radio-telemetry probe (HD-X10; Data Sciences International) as previously described (66). Briefly, under isoflurane anesthesia, a neck incision was made, and the tip of catheter was inserted into the left carotid artery and secured by ligation with 6-0 silk suture. The transducer/telemetry transmitter was placed on the dorsum in a subcutaneous location. Continuous blood pressure, heart rate, and body temperature data were averaged for each minute and transmitted telemetrically. To reduce variability, we averaged consecutive 1-hour time windows starting immediately after CLP surgery.

### Renal hypoxia staining in vivo

Renal hypoxia was assessed by pimonidazole immunohistochemistry. Pimonidazole (Hypoxyprobe-1 in Hypoxyprobe-1 Omni Kit; Hypoxyprobe Inc., Burlington, MA, 60 mg/kg) was administrated into mice intraperitoneally at 2h before euthanasia. Kidneys were fixed with 10% formalin and processed for paraffin embedding, sectioning, peroxidase quenching and blocking as described previously(24). The sections were sequentially incubated with rabbit anti- pimonidazole antibody (PAb2627; Hypoxyprobe Inc., Burlington, MA), and subsequently polyclonal goat anti-rabbit Immunoglobulins conjugated with HRP (Agilent Dako, Santa Clara, CA), following the manufacturer instructions.

Renal hypoxia was evaluated by semiquantitative measurements of pimonidazole staining on tubules of the cortex and the outer stripe of the OSOM. The degree of pimonidazole staining was estimated at 400x magnification using more than five randomly selected fields for each animal, and scored according to the intensity and the extent of positive cells: 0, no staining; 1, minimal, 2; moderate, 3; severe staining.

### Clinical chemistry markers and cytokines

An autoanalyzer (Hitachi 917; Boehringer Mannheim, Indianapolis, IN) was used to measure serum BUN, lactate dehydrogenase, alkaline phosphatase, aspartate transaminase. Serum creatinine was measured by HPLC(67). Serum IL-6, IL-10, IL-17, and TNFα were measured by ELISA (R & D Systems, Minneapolis, MN).

### qPCR of mtDNA

Plasma and urinary DNA of samples were extracted with the QIAamp DNA Mini and Blood Mini kit (Qiagen, Germantown, MD). Taqman primers (forward: 5’-TCATGTCGGACGAGGCTTAT-3’, reverse: 5’-CTCATGGAAGGACGTAGCCT-3’) and a probe (5’-ACTGTTCGCAGTCATAGCCACAGCA-3’) with a unique 5′fluorescent reporter dye (FAM) and 3 fluorescent quencher dye (TAMRA) were designed using mouse mitochondrial genome sequences and synthesized by Genscript (Piscataway, NJ). For absolute quantification using qPCR, a mitochondrial DNA standard was prepared by PCR of liver mitochondrial DNA with these primers, purified using a QIAquick PCR Purification Kit (Qiagen, Germantown, MD), and serially diluted. The concentration of the mitochondrial DNA standard was measured by Qubit 1XdsDNA HS Assay Kit with Qubit 4 Fluorometer (Thermo Fisher Scientific, Waltham, MA). For the experiment described in **Supplemental Figure 4**, CD-1 mice were subjected to either sham or CLP surgery, then 18 hours later, livers were removed, and liver mitochondria were isolated from with Mitochondria Isolation Kit for Tissue (Abcam, Cambridge, MA).Liver mitochondrial DNA was then extracted from mitochondria using DNeasy Blood & Tissue kit (Qiagen, Germantown, MD). With the standard curves of the targeted mtDNA, plasma and urinary mtDNA in samples was quantified by the following thermal profile used with QuantStudio™ 6 Flex System (Applied Biosystems, Carlsbad, CA, consisting of an initial cycle of 2 minutes at 95 °C, followed by 40 cycles of 5 seconds at 95 °C and 10 seconds at 54 °C. The plasma concentration of mtDNA is expressed as copies per microliter; urine mtDNA is expressed as copies per gram creatinine, as measured by Creatinine Companion (Ethos Biosciences, Newtown Square, PA).

### Measuring mitochondrial respiratory function

A Cell Mito Stress Test (Agilent, Santa Clara, CA) was performed following the manufacturer’s protocol on a Seahorse XFe96 extracellular flux analyzer (Agilent). TECs were plated on a sterile XFe96 well plate in duplicate at 2 × 10^4^ total cells per well and serially stimulated in the following sequence: (A) 1μM oligomycin; (B) 0-100 μM carbonyl cyanide-4-(trifluoromethoxy) phenylhydrazone (FCCP) or BAM15; (C) 2 μM rotenone/antimycin A; (D) 4μg/ml Hoechst 33342. The plate was returned to the Cytation 5 (Bioteck, Winooski, VT) for a fluorescence scan and cell count. Normalization, calculation, and visualization of respiratory parameters including basal oxygen consumption rate (OCR) and maximum respiration were performed by Wave software (Agilent, Santa Clara, CA).

### Live cell imaging of mtROS

Mouse primary PTECs were plated onto a chambered cover glass and incubated in phenol-red-free low glucose (5.5mM) medium for one day and stained with 200nM MitoSOXTM Red and 2 μg/ml Hoechst 33343 (Thermo Fisher Scientific, Waltham, MA) for 30 min at 37°C. PTECs were washed with Hanks’ balanced salt solution (HBSS) with Ca and Mg, and were incubated with ProLong Live Antifade Reagent (Thermo Fisher Scientific, Waltham, MA). Serum was collected from mice at 18 hours after CLP or sham surgery and was filtered through a 0.22μm pore filter. Renal Epithelial Cell Growth Medium 2 (PromoCell, Heidelberg, Germany) was supplemented with a) 10% serum from mice subjected to either CLP or sham surgery (collected 18 hours post-surgery) and b) 10 or 20μM BAM15 or the equivalent volume of vehicle. PTECs were incubated in these media.

Images of cultured cells were acquired by confocal microscopy with incubation system (Zeiss LSM780, Zeiss, Oberkochen, German) at 0, 6, 12 and 24 hours after serum addition. Isolated mtDNA (1ng) from liver of Sham or CLP mice also were incubated with mouse primary PTECs. The images were acquired at 3 hours after adding mtDNA. The fluorescence of MitoSOX™ Red and Hoechst 33343 were imaged at 488 nm and 405 nm excitation and using 562nm-666nm and 413mm-490nm emission ranges. All imaging parameters remained the same during data acquisition, and images were processed identically and exported using Zen Zeiss software. The relative fluorescence units of individual cells were quantified using Fiji/ ImageJ software. Corrected total cell fluorescence units were determined using the following formula: integrated density of selected cell − (area of selected cell × mean integrated density of background readings).

### Bacterial counts

Blood specimens and peritoneal exudate, obtained after injection of 0.9% saline (2 ml) into the peritoneal cavity, were collected under sterile conditions at 18 hours after CLP or Sham surgery. Serial dilutions were plated on 5% sheep blood agar (BD, Franklin Lakes, NJ) and incubated at 37°C. The number of colonies was counted after cultured for 24 hours at 37°C.

Cecal material was collected from healthy CD1-mice, and diluted with 10ml of PBS as bacterial suspension of CLP. Total 50ul of bacterial suspension treated with 0, 1, 5, 10, 20 and 50μM BAM15 was applied to BBL trypticase soy agar with 5% sheep blood (TSAⅡ) (BD, Franklin Lakes, NJ). The number of colonies was counted after culture for 24 hours at 37°C.

### Western-blotting analysis

Total protein was extracted from whole kidney based using a bead based Precellys 24 tissue homogenizer (Bertin Technologies, Montigny-le-Bretonneux, France) with T-PER Tissue Protein Extraction Reagent (Thermo Fisher Scientific, Waltham, MA) treated with Halt Phosphatase inhibitor Cocktail (Thermo Fisher Scientific, Waltham, MA) and Complete Mini Protease Inhibitor Cocktail (Millipore Sigma, Burlington, MA). Lysate protein concentrations were measured by BCA assay and adjusted to be the same. Protein lysates were denatured by mixing with an equal volume of 2X Laemmli sample buffer with 0.2 M DTT and heated for 10 min at 95°. Samples were resolved by SDS-PAGE and transferred to nitrocellulose membranes using an iBlot dry blotting system (Invitrogen, Carlsbad, CA). Membranes were blocked for 30 min with blocking buffer for fluorescent Western blotting (Rockland Immunochemicals, Inc., Pottstown, PA) and incubated with the primary antibody overnight at 4°C. Membranes were washed and incubated with secondary antibody for 1 hour at room temperature. Primary antibodies included anti-AMPKα mouse antibody (1:500, #2793), Phospho-AMPKα rabbit antibody (1:500, #2535), anti-Sirt1 rabbit antibody (1:1000, #9475, all from Cell signaling technology, Danvers, MA) anti-PGC1α rabbit antibody (1:1000, NBP1-04676, Novus Biologicals, Centennial, CO) anti-TFAM rabbit antibody (1:1000, SAB 1401383, Sigma-Aldrich, Saint Louis, MO), anti-β-actin mouse antibody as a control (1:5000, #3700, Cell Signaling Technology, Danvers, MA). Labeled secondary antibodies included IRDye 680LT Goat anti-rabbit IgG secondary antibody (1:4000) and IRDye 800CW donkey anti-mouse IgG secondary antibody (1:3000, both from LI-COR Biosciences, Lincoln, NE). Western blot membranes were imaged by two-color Western blot detection with infrared fluorescence with a LI-COR Odyssey infrared scanner (LI-COR Biosciences, Lincoln, NE). Band intensities were analyzed using Odyssey software.

### Quantitation of NAD^+^/NADH in kidney

Fresh kidneys were harvested from mice and 25 mg of the cortex was used for assay. Kidneys were homogenized by a bead based Precellys 24 tissue homogenizer (Bertin Technologies, Montigny-le-Bretonneux, France). NAD + and NADH in kidney lysate were measured following the procedure of BioAssay Systems’ EnzyChrom NAD+/NADH assay kit (BioAssay Systems, Hayward, CA). The optical density (OD) values for time “zero” and OD15 after a 15-min incubation were obtained at 565 nm.

### Statistics

Data are presented as mean ± SEM. Student’s t-test was applied when comparing two groups for which both datasets had a normal distribution. Comparison of three or more groups was assessed by one-way ANOVA, and the influence of two different independent variables (i.e., time point and group) on an outcome was assessed by two-way ANOVA when data was normally distributed. Mixed-effects analysis was performed instead of two-way ANOVA to handle missing data. Dunn’s multiple comparisons test was used with Kruskal-Wallis test when the data were not normally distributed. As a post-hoc test, Tukey’s multiple comparisons test was used post-hoc for both one-way and two-way ANOVA analyses. Sidak’s multiple comparisons test was used for mixed-effects analysis. The prediction ability for mortality data was assessed using receiver operating characteristic (ROC) curve analysis and log-rank test. All statistical analyses were performed using Prism version 8.4.3 (GraphPad Software, La Jolla, CA).

## Author contributions

N.T., P.S.Y. and R.A.S. conceived and designed research; N.T., T.T., T.Y., and X.H. performed experiments; N.T. analyzed data; N.T., T.T., P.S.Y., and R.A.S. interpreted results of experiments; N.T. prepared figures; N.T. drafted manuscript; N.T., P.S.Y., and R.A.S. edited and revised manuscript; N.T., T.T., T.Y., X.H. P.S.Y. P.S.Y., and R.A.S. approved final version of manuscript.

## Acknowledgments

This research was supported by the Intramural Research Program of the National Institutes of Health, National Institute of Diabetes and Digestive and Kidney Diseases (NIDDK). Naoko Tsuji is also supported in part by a Japan Society for the Promotion of Science Research Fellowship for Japanese Biomedical and Behavioral Researchers at the National Institutes of Health (No. 2-72005). The authors thank Jeff M. Reece for setup and advice on confocal microscopy and Jeffrey B Kopp for detailed editing of the manuscript. The authors have declared that no conflict of interest exists.

---

**Supplemental Figure 1. BAM15 treatment improves mortality and AKI in septic female mice.** (A) Kaplan-Meier curves of Sham or CLP female mice treated with vehicle (at 0 hours) and Sham or CLP treated with BAM15 (5mg/kg, at 0 hours) for 7 days. Sham+ Vehicle/ BAM15: n=4 each, CLP+ Vehicle/ BAM15: n=20 each. Log-rank test. *p<0.05, CLP + Vehicle vs other groups. (B-C) BUN (B), AST, ALT, LDH, Amylase, CK (C) by biochemical examination at 18hrs after Sham or CLP female mice treated with vehicle (at 0 hours) or BAM15 (5mg/kg, at 0 hours). All bars show mean ± SEM of each group (Sham + Vehicle: n=6, CLP + Vehicle: n=12, Sham + BAM15: n=6, CLP + BAM15: n=12). Dunn’s multiple comparisons test following Kruskal Wallis test. *: vs Sham + Vehicle, p<0.05. †: vs Sham + BAM15, p<0.05. ‡: vs CLP + Vehicle, p<0.05. (D-F) Time course of plasma mtDNA level (D) and urine mtDNA (E) and urine mtDNA adjusted to the creatinine excretion (F) at 18 hours after CLP (n=12 each) mice treated with vehicle (at 0 hours) or BAM15 (5mg/kg, at 0 hours). The data represent mean ± SEM. Analysis between groups at each time point was performed with Sidak’s multiple comparisons test following mixed-effects analysis. *: p<0.05.

**Supplemental Figure 2. BAM15 does not kill bacteria.** (A) Bacterial count in Blood and fluid of abdominal cavity at 18 hours after Sham (n=5 each) or CLP (n=4 each) mice treated with vehicle (at 0 hours) or BAM15 (5mg/kg, at 0 hours). (B) Bacterial count in suspension of cecum material treated with 0, 1, 5, 10, 20, and 50 μM BAM15 (n=3~4). All graph represent mean±SEM. Tukey’s multiple comparisons test following one-way ANOVA test. * vs Sham + Vehicle, p<0.05. † vs Sham + BAM15, p<0.05. ‡ vs CLP + Vehicle, p<0.05.

**Supplemental Figure 3. Biological activity of BAM15 on mitochondrial respiratory chain as an uncoupler.** (A-D) OCR and Maximum respiration of FCCP (A, B) and BAM15 (C, D) with 4 biological replicates per each concentration (0, 1, 2, 5, 10, 20, 50, and 100μM) and the concentration dependency on the maximum respiration (E). Tukey’s multiple comparisons test following one-way ANOVA test (B, D) and Sidak’s multiple comparisons test following two-way ANOVA test (E). *: p<0.05.

**Supplemental Figure 4. Generation of mtROS in kidney tubule cells by sepsis-derived mtDNA.** (A) MitoSOX-Red intensity in mouse primary tubular cells treated with mtDNA (1 ng) purified from liver of CLP or Sham mice. Each dot shows MitoSOX-Red intensity of a cell from 5 randomized images (magnification x400) per well on 2 biological replicates per each condition. Tukey’s multiple comparison test following one-way ANOVA test. *: p<0.05.

## References

1. Rudd KE, Johnson SC, Agesa KM, Shackelford KA, Tsoi D, Kievlan DR, et al. Global, regional, and national sepsis incidence and mortality, 1990-2017: analysis for the Global Burden of Disease Study. Lancet. 2020;395(10219):200–11.

2. Murugan R, Karajala-Subramanyam V, Lee M, Yende S, Kong L, Carter M, et al. Acute kidney injury in non-severe pneumonia is associated with an increased immune response and lower survival. Kidney Int. 2010;77(6):527–35.

3. Hoste EA, Bagshaw SM, Bellomo R, Cely CM, Colman R, Cruz DN, et al. Epidemiology of acute kidney injury in critically ill patients: the multinational AKI-EPI study. Intensive Care Med. 2015;41(8):1411–23.

4. Kimmel PL, Jefferson N, Norton JM, and Star RA. How Community Engagement Is Enhancing NIDDK Research. Clin J Am Soc Nephrol. 2019;14(5):768–70.

5. Reddy K, Sinha P, O’Kane CM, Gordon AC, Calfee CS, and McAuley DF. Subphenotypes in critical care: translation into clinical practice. The Lancet Respiratory Medicine. 2020;8(6):631–43.

6. Papadopoulos N, Kinzler KW, and Vogelstein B. The role of companion diagnostics in the development and use of mutation-targeted cancer therapies. Nat Biotechnol. 2006;24(8):985–95.

7. Tsuji N, and Agbor-Enoh S. Cell-free DNA beyond a biomarker for rejection: Biological trigger of tissue injury and potential therapeutics. J Heart Lung Transplant. 2021.

8. Tsuji N, Tsuji T, Ohashi N, Kato A, Fujigaki Y, and Yasuda H. Role of Mitochondrial DNA in Septic AKI via Toll-Like Receptor 9. J Am Soc Nephrol. 2016;27(7):2009–20.

9. Krychtiuk KA, Ruhittel S, Hohensinner PJ, Koller L, Kaun C, Lenz M, et al. Mitochondrial DNA and Toll-Like Receptor-9 Are Associated With Mortality in Critically Ill Patients. Crit Care Med. 2015;43(12):2633–41.

10. Scozzi D, Cano M, Ma L, Zhou D, Zhu JH, O’Halloran JA, et al. Circulating mitochondrial DNA is an early indicator of severe illness and mortality from COVID-19. JCI Insight. 2021;6(4).

11. Andargie TE, Tsuji N, Seifuddin F, Jang MK, Yuen PST, Kong H, et al. Cell-free DNA maps COVID-19 tissue injury and risk of death and can cause tissue injury. JCI Insight. 2021;6(7).

12. Zuo Y, Yalavarthi S, Shi H, Gockman K, Zuo M, Madison JA, et al. Neutrophil extracellular traps in COVID-19. JCI Insight. 2020;5(11).

13. Kenwood BM, Weaver JL, Bajwa A, Poon IK, Byrne FL, Murrow BA, et al. Identification of a novel mitochondrial uncoupler that does not depolarize the plasma membrane. Mol Metab. 2014;3(2):114–23.

14. Ou J, Ball JM, Luan Y, Zhao T, Miyagishima KJ, Xu Y, et al. iPSCs from a Hibernator Provide a Platform for Studying Cold Adaptation and Its Potential Medical Applications. Cell. 2018;173(4):851–63 e16.

15. Nishikawa T, Edelstein D, Du XL, Yamagishi S, Matsumura T, Kaneda Y, et al. Normalizing mitochondrial superoxide production blocks three pathways of hyperglycaemic damage. Nature. 2000;404(6779):787–90.

16. Tahara EB, Navarete FD, and Kowaltowski AJ. Tissue-, substrate-, and site-specific characteristics of mitochondrial reactive oxygen species generation. Free Radic Biol Med. 2009;46(9):1283–97.

17. Childress ES, Alexopoulos SJ, Hoehn KL, and Santos WL. Small Molecule Mitochondrial Uncouplers and Their Therapeutic Potential. J Med Chem. 2018;61(11):4641–55.

18. Axelrod CL, King WT, Davuluri G, Noland RC, Hall J, Hull M, et al. BAM15-mediated mitochondrial uncoupling protects against obesity and improves glycemic control. EMBO Mol Med. 2020;12(7):e12088.

19. Tao H, Zhang Y, Zeng X, Shulman GI, and Jin S. Niclosamide ethanolamine-induced mild mitochondrial uncoupling improves diabetic symptoms in mice. Nat Med. 2014;20(11):1263–9.

20. Tran M, Tam D, Bardia A, Bhasin M, Rowe GC, Kher A, et al. PGC-1alpha promotes recovery after acute kidney injury during systemic inflammation in mice. J Clin Invest. 2011;121(10):4003–14.

21. Lynch MR, Tran MT, Ralto KM, Zsengeller ZK, Raman V, Bhasin SS, et al. TFEB-driven lysosomal biogenesis is pivotal for PGC1alpha-dependent renal stress resistance. JCI Insight. 2019;5.

22. Miyaji T, Hu X, Yuen PS, Muramatsu Y, Iyer S, Hewitt SM, et al. Ethyl pyruvate decreases sepsis-induced acute renal failure and multiple organ damage in aged mice. Kidney Int. 2003;64(5):1620–31.

23. Doi K, Leelahavanichkul A, Hu X, Sidransky KL, Zhou H, Qin Y, et al. Pre-existing renal disease promotes sepsis-induced acute kidney injury and worsens outcome. Kidney Int. 2008;74(8):1017–25.

24. Yasuda H, Yuen PS, Hu X, Zhou H, and Star RA. Simvastatin improves sepsis-induced mortality and acute kidney injury via renal vascular effects. Kidney Int. 2006;69(9):1535–42.

25. Kracke F, Vassilev I, and Kromer JO. Microbial electron transport and energy conservation - the foundation for optimizing bioelectrochemical systems. Front Microbiol. 2015;6:575.

26. Harrington JS, Huh JW, Schenck EJ, Nakahira K, Siempos, II, and Choi AMK. Circulating Mitochondrial DNA as Predictor of Mortality in Critically Ill Patients: A Systematic Review of Clinical Studies. Chest. 2019;156(6):1120–36.

27. Dennis JM, and Witting PK. Protective Role for Antioxidants in Acute Kidney Disease. Nutrients. 2017;9(7).

28. Seija M, Baccino C, Nin N, Sanchez-Rodriguez C, Granados R, Ferruelo A, et al. Role of peroxynitrite in sepsis-induced acute kidney injury in an experimental model of sepsis in rats. Shock. 2012;38(4):403–10.

29. Sun J, Zhang J, Tian J, Virzi GM, Digvijay K, Cueto L, et al. Mitochondria in Sepsis-Induced AKI. J Am Soc Nephrol. 2019;30(7):1151–61.

30. Gomez H, Kellum JA, and Ronco C. Metabolic reprogramming and tolerance during sepsis-induced AKI. Nat Rev Nephrol. 2017;13(3):143–51.

31. Ralto KM, Rhee EP, and Parikh SM. NAD(+) homeostasis in renal health and disease. Nat Rev Nephrol. 2020;16(2):99–111.

32. Kang I, Chu CT, and Kaufman BA. The mitochondrial transcription factor TFAM in neurodegeneration: emerging evidence and mechanisms. FEBS Lett. 2018;592(5):793–811.

33. Guo Y, Ni J, Chen S, Bai M, Lin J, Ding G, et al. MicroRNA-709 Mediates Acute Tubular Injury through Effects on Mitochondrial Function. J Am Soc Nephrol. 2018;29(2):449–61.

34. Venet F, and Monneret G. Advances in the understanding and treatment of sepsis-induced immunosuppression. Nat Rev Nephrol. 2018;14(2):121–37.

35. Schaefer L. Complexity of danger: the diverse nature of damage-associated molecular patterns. J Biol Chem. 2014;289(51):35237–45.

36. Yasuda H, Leelahavanichkul A, Tsunoda S, Dear JW, Takahashi Y, Ito S, et al. Chloroquine and inhibition of Toll-like receptor 9 protect from sepsis-induced acute kidney injury. Am J Physiol Renal Physiol. 2008;294(5):F1050–8.

37. West AP, and Shadel GS. Mitochondrial DNA in innate immune responses and inflammatory pathology. Nat Rev Immunol. 2017;17(6):363–75.

38. Unsinger J, McGlynn M, Kasten KR, Hoekzema AS, Watanabe E, Muenzer JT, et al. IL-7 promotes T cell viability, trafficking, and functionality and improves survival in sepsis. J Immunol. 2010;184(7):3768–79.

39. Brahmamdam P, Inoue S, Unsinger J, Chang KC, McDunn JE, and Hotchkiss RS. Delayed administration of anti-PD-1 antibody reverses immune dysfunction and improves survival during sepsis. J Leukoc Biol. 2010;88(2):233–40.

40. Camerota AJ, Creasey AA, Patla V, Larkin VA, and Fink MP. Delayed treatment with recombinant human tissue factor pathway inhibitor improves survival in rabbits with gram-negative peritonitis. J Infect Dis. 1998;177(3):668–76.

41. Holthoff JH, Wang Z, Seely KA, Gokden N, and Mayeux PR. Resveratrol improves renal microcirculation, protects the tubular epithelium, and prolongs survival in a mouse model of sepsis-induced acute kidney injury. Kidney Int. 2012;81(4):370–8.

42. Feng X, Zhu W, Schurig-Briccio LA, Lindert S, Shoen C, Hitchings R, et al. Antiinfectives targeting enzymes and the proton motive force. Proc Natl Acad Sci U S A. 2015;112(51):E7073–82.

43. Tang M, Luo Z, Wu Y, Zhuang J, Li K, Hu D, et al. BAM15 attenuates transportation-induced apoptosis in iPS-differentiated retinal tissue. Stem Cell Res Ther. 2019;10(1):64.

44. Zhao M, Wang Y, Li L, Liu S, Wang C, Yuan Y, et al. Mitochondrial ROS promote mitochondrial dysfunction and inflammation in ischemic acute kidney injury by disrupting TFAM-mediated mtDNA maintenance. Theranostics. 2021;11(4):1845–63.

45. Wang Y, Zhu J, Liu Z, Shu S, Fu Y, Liu Y, et al. The PINK1/PARK2/optineurin pathway of mitophagy is activated for protection in septic acute kidney injury. Redox Biol. 2021;38:101767.

46. Hu Q, Ren J, Ren H, Wu J, Wu X, Liu S, et al. Urinary Mitochondrial DNA Identifies Renal Dysfunction and Mitochondrial Damage in Sepsis-Induced Acute Kidney Injury. Oxid Med Cell Longev. 2018;2018:8074936.

47. Whitaker RM, Stallons LJ, Kneff JE, Alge JL, Harmon JL, Rahn JJ, et al. Urinary mitochondrial DNA is a biomarker of mitochondrial disruption and renal dysfunction in acute kidney injury. Kidney Int. 2015;88(6):1336–44.

48. Hu Q, Ren J, Wu J, Li G, Wu X, Liu S, et al. Urinary Mitochondrial DNA Levels Identify Acute Kidney Injury in Surgical Critical Illness Patients. Shock. 2017;48(1):11–7.

49. Timmermans K, Kox M, Scheffer GJ, and Pickkers P. Plasma Nuclear and Mitochondrial DNA Levels, and Markers of Inflammation, Shock, and Organ Damage in Patients with Septic Shock. Shock. 2016;45(6):607–12.

50. van der Slikke EC, Star BS, van Meurs M, Henning RH, Moser J, and Bouma HR. Sepsis is associated with mitochondrial DNA damage and a reduced mitochondrial mass in the kidney of patients with sepsis-AKI. Crit Care. 2021;25(1):36.

51. Collier JB, and Schnellmann RG. Extracellular Signal-Regulated Kinase 1/2 Regulates Mouse Kidney Injury Molecule-1 Expression Physiologically and Following Ischemic and Septic Renal Injury. J Pharmacol Exp Ther. 2017;363(3):419–27.

52. Yamamoto T, Noiri E, Ono Y, Doi K, Negishi K, Kamijo A, et al. Renal L-type fatty acid--binding protein in acute ischemic injury. J Am Soc Nephrol. 2007;18(11):2894–902.

53. Chambers JM, and Wingert RA. PGC-1alpha in Disease: Recent Renal Insights into a Versatile Metabolic Regulator. Cells. 2020;9(10).

54. Price NL, Gomes AP, Ling AJ, Duarte FV, Martin-Montalvo A, North BJ, et al. SIRT1 is required for AMPK activation and the beneficial effects of resveratrol on mitochondrial function. Cell Metab. 2012;15(5):675–90.

55. Feingold KR, Wang Y, Moser A, Shigenaga JK, and Grunfeld C. LPS decreases fatty acid oxidation and nuclear hormone receptors in the kidney. J Lipid Res. 2008;49(10):2179–87.

56. Escobar DA, Botero-Quintero AM, Kautza BC, Luciano J, Loughran P, Darwiche S, et al. Adenosine monophosphate-activated protein kinase activation protects against sepsis-induced organ injury and inflammation. J Surg Res. 2015;194(1):262–72.

57. Opal SM, Ellis JL, Suri V, Freudenberg JM, Vlasuk GP, Li Y, et al. Pharmacological Sirt1 Activation Improves Mortality and Markedly Alters Transcriptional Profiles That Accompany Experimental Sepsis. Shock. 2016;45(4):411–8.

58. Tai Y, Li L, Peng X, Zhu J, Mao X, Qin N, et al. Mitochondrial uncoupler BAM15 inhibits artery constriction and potently activates AMPK in vascular smooth muscle cells. Acta Pharm Sin B. 2018;8(6):909–18.

59. Gureev AP, Shaforostova EA, and Popov VN. Regulation of Mitochondrial Biogenesis as a Way for Active Longevity: Interaction Between the Nrf2 and PGC-1alpha Signaling Pathways. Front Genet. 2019;10:435.

60. West AP, Khoury-Hanold W, Staron M, Tal MC, Pineda CM, Lang SM, et al. Mitochondrial DNA stress primes the antiviral innate immune response. Nature. 2015;520(7548):553–7.

61. Dear JW, Yasuda H, Hu X, Hieny S, Yuen PS, Hewitt SM, et al. Sepsis-induced organ failure is mediated by different pathways in the kidney and liver: acute renal failure is dependent on MyD88 but not renal cell apoptosis. Kidney Int. 2006;69(5):832–6.

62. Osuchowski MF, Ayala A, Bahrami S, Bauer M, Boros M, Cavaillon JM, et al. Minimum Quality Threshold in Pre-Clinical Sepsis Studies (MQTiPSS): an international expert consensus initiative for improvement of animal modeling in sepsis. Infection. 2018;46(5):687–91.

63. Zafrani, L, Gerotziafas, G, Byrnes, C, Hu, X, Perez, J, Levi, C, Placier, S, Letavernier, E, Leelahavanichkul, A, Haymann, JP, Elalamy, I, Miller, JL, Star, RA, Yuen, PST, and Baud, L. Calpastatin controls polymicrobial sepsis by limiting procoagulant microparticle release. Am J Resp Crit Care Med. 2012; 185(7):744–55.

64. Iwakura T, Zhao Z, Marschner JA, Devarapu SK, Yasuda H, and Anders HJ. Dipeptidyl peptidase-4 inhibitor teneligliptin accelerates recovery from cisplatin-induced acute kidney injury by attenuating inflammation and promoting tubular regeneration. Nephrol Dial Transplant. 2019;34(10):1669–80.

65. Terryn S, Jouret F, Vandenabeele F, Smolders I, Moreels M, Devuyst O, et al. A primary culture of mouse proximal tubular cells, established on collagen-coated membranes. Am J Physiol Renal Physiol. 2007;293(2):F476–85.

66. Street JM, Koritzinsky EH, Bellomo TR, Hu X, Yuen PST, and Star RA. The role of adenosine 1a receptor signaling on GFR early after the induction of sepsis. Am J Physiol Renal Physiol. 2018;314(5):F788–F97.

67. Yuen PS, Dunn SR, Miyaji T, Yasuda H, Sharma K, and Star RA. A simplified method for HPLC determination of creatinine in mouse serum. Am J Physiol Renal Physiol. 2004;286(6):F1116–9.

