## Supplemental Figures for "BAM15 treats mouse sepsis and sepsis-AKI, linking circulating mitochondrial DNA and tubule reactive oxygen species"

Supplemental figure 1

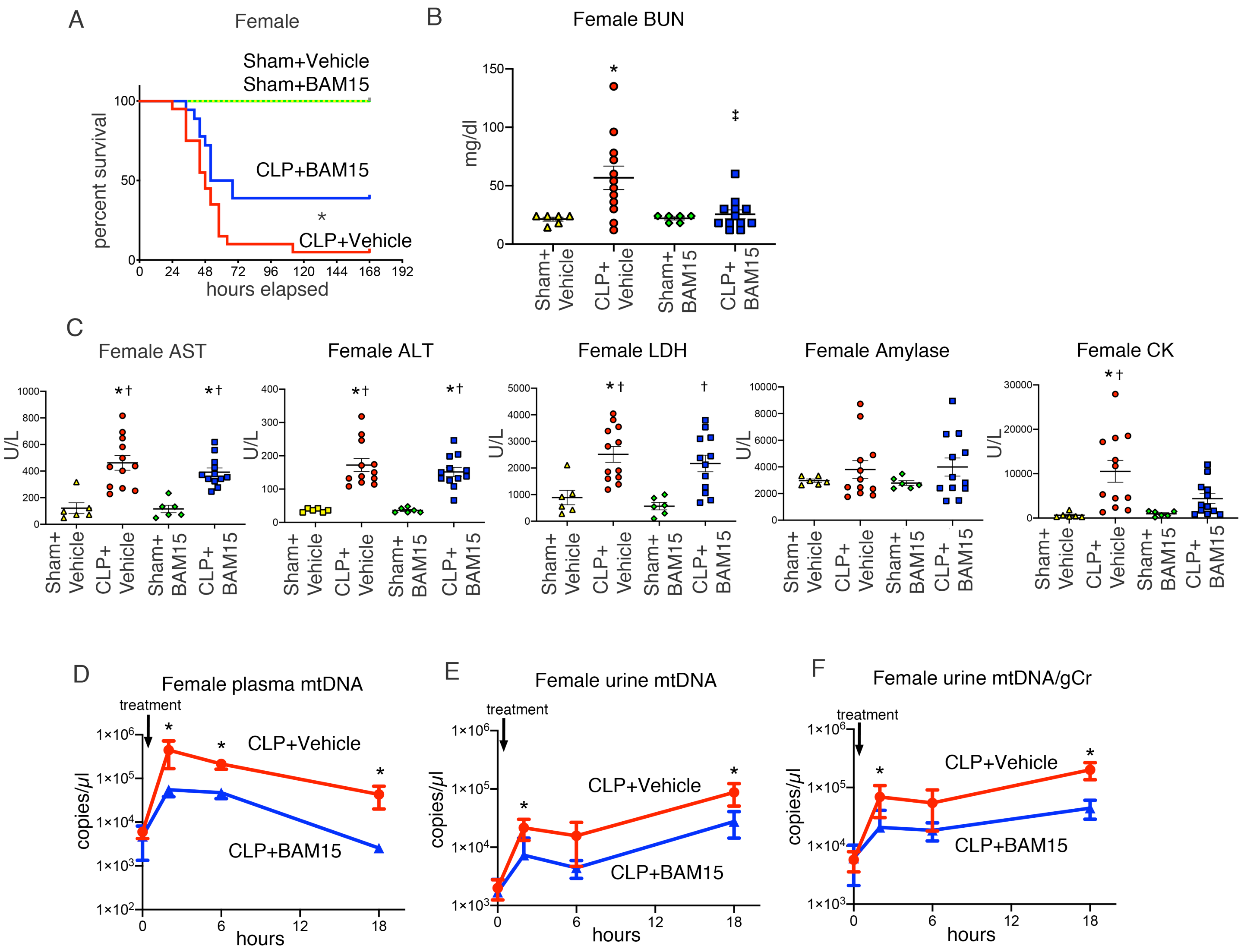


**Supplemental Figure 1. BAM15 treatment improves mortality and AKI in septic female mice.** (A) Kaplan-Meier curves of Sham or CLP female mice treated with vehicle (at 0 hours) and Sham or CLP treated with BAM15 (5mg/kg, at 0 hours) for 7 days. Sham+ Vehicle/ BAM15: n=4 each, CLP+ Vehicle/ BAM15: n=20 each. Log-rank test. *p<0.05, CLP + Vehicle vs other groups. (B-C) BUN (B), AST, ALT, LDH, Amylase, CK (C) by biochemical examination at 18hrs after Sham or CLP female mice treated with vehicle (at 0 hours) or BAM15 (5mg/kg, at 0 hours). All bars show mean ± SEM of each group (Sham + Vehicle: n=6, CLP + Vehicle: n=12, Sham + BAM15: n=6, CLP + BAM15: n=12). Dunn's multiple comparisons test following Kruskal Wallis test. *: vs Sham + Vehicle, p<0.05. †: vs Sham + BAM15, p<0.05. ‡: vs CLP + Vehicle, p<0.05. (D-F) Time course of plasma mtDNA level (D) and urine mtDNA (E) and urine mtDNA adjusted to the creatinine excretion (F) at 18 hours after CLP (n=12 each) mice treated with vehicle (at 0 hours) or BAM15 (5mg/kg, at 0 hours). The data represent mean ± SEM. Analysis between groups at each time point was performed with Sidak's multiple comparisons test following mixed-effects analysis. *: p<0.05.

Supplemental figure 2

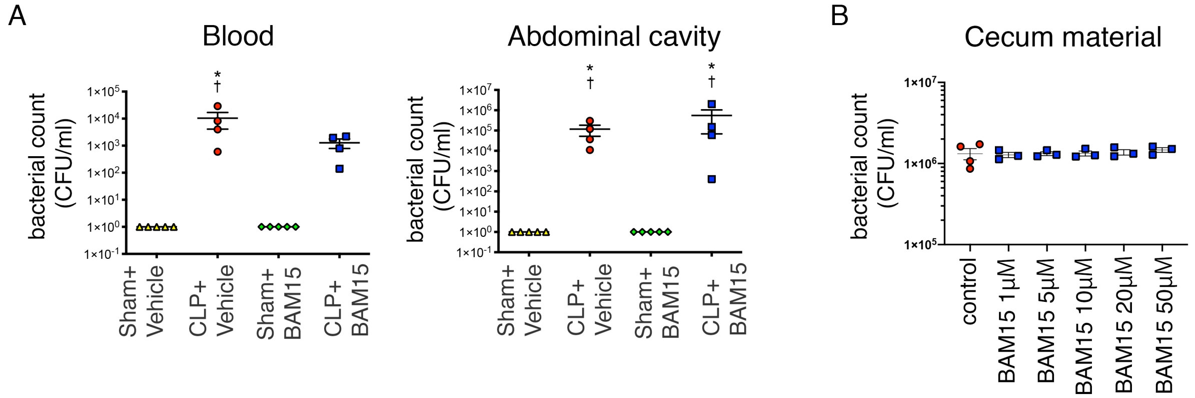


**Supplemental Figure 2. BAM15 does not kill bacteria.** (A) Bacterial count in Blood and fluid of abdominal cavity at 18 hours after Sham (n=5 each) or CLP (n=4 each) mice treated with vehicle (at 0 hours) or BAM15 (5mg/kg, at 0 hours). (B) Bacterial count in suspension of cecum material treated with 0, 1, 5, 10, 20, and 50 $\mu$M BAM15 (n=3~4). All graph represent mean±SEM. Tukey's multiple comparisons test following one-way ANOVA test. * vs Sham + Vehicle, $p<0.05$. † vs Sham + BAM15, $p<0.05$. ‡ vs CLP + Vehicle, $p<0.05$.

Supplemental figure 3

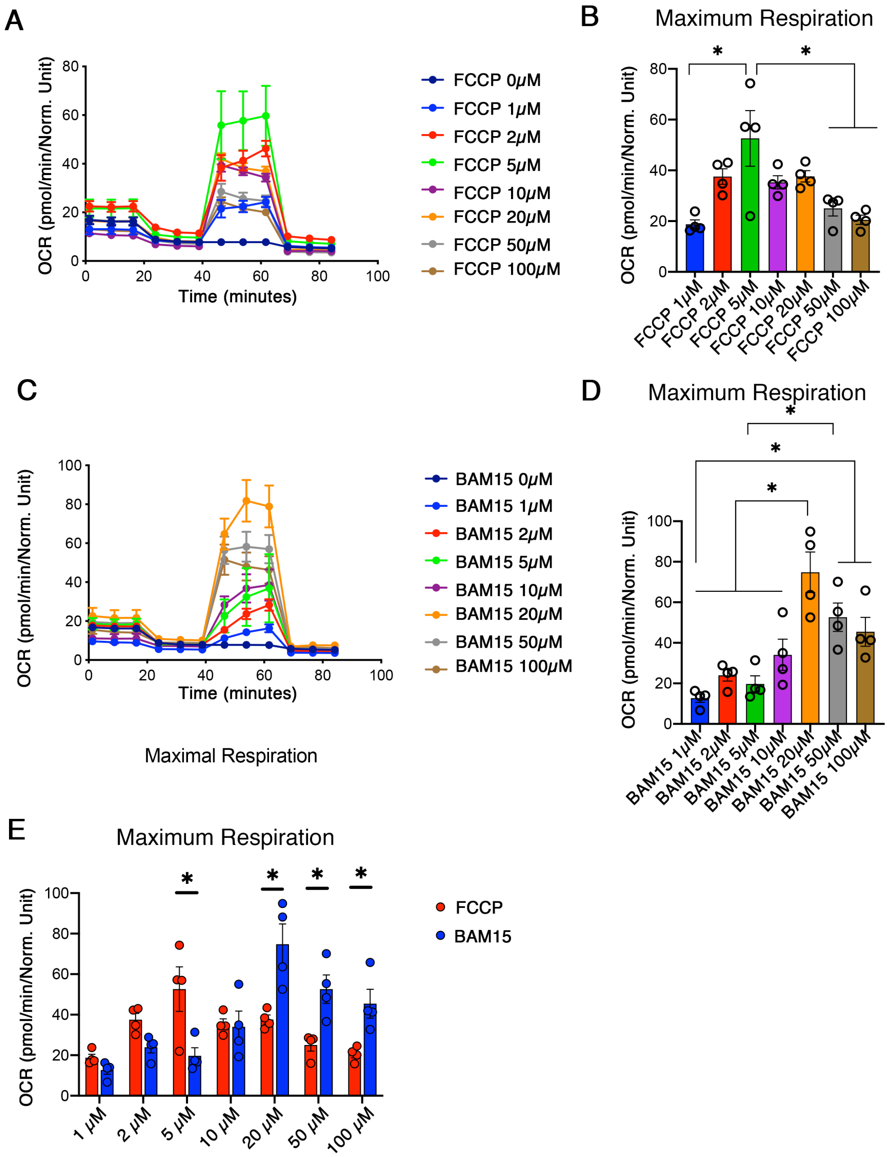


**Supplemental Figure 3. Biological activity of BAM15 on mitochondrial respiratory chain as an uncoupler.** (A-D) OCR and Maximum respiration of FCCP (A, B) and BAM15 (C, D) with 4 biological replicates per each concentration (0, 1, 2, 5, 10, 20, 50, and 100μM) and the concentration dependency on the maximum respiration (E). Tukey's multiple comparisons test following one-way ANOVA test (B, D) and Sidak's multiple comparisons test following two-way ANOVA test (E). *: $p<0.05$.

Supplemental figure 4

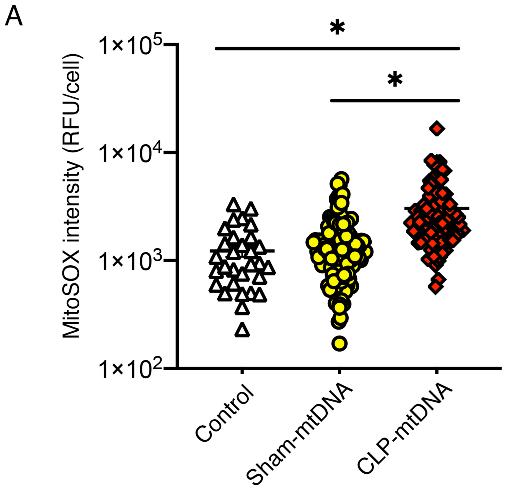


**Supplemental Figure 4. Generation of mtROS in kidney tubule cells by sepsis-derived mtDNA.** (A) MitoSOX-Red intensity in mouse primary tubular cells treated with mtDNA (1 ng) purified from liver of CLP or Sham mice. Each dot shows MitoSOX-Red intensity of a cell from 5 randomized images (magnification x400) per well on 2 biological replicates per each condition. Tukey's multiple comparison test following one-way ANOVA test. *: $p<0.05$.
